# Azimuthal Segment Imaging in cryo-STEM Tomography

**DOI:** 10.1101/2025.11.26.690630

**Authors:** Peter Kirchweger, Shahar Seifer, Sharon Grayer Wolf, Neta Varsano, Bettina Zens, Florian K.M. Schur, Michael Elbaum

## Abstract

Cryo-electron microscopy is transitioning from investigation of isolated macromolecules to in situ studies bridging the realms of structural and cellular biology. Newly available detector technologies enable unconventional contrast modes with particular advantages. Here we demonstrate application of quadrant diode detectors to visualize a range of biological specimens by cryo-Scanning Transmission Electron Tomography (cryo-STET). Theoretically, we decompose coherent contrast by parallax analysis to isolate phase and amplitude contributions in specimens too thick for energy-filtered TEM. We thereby expand the cryo-STEM toolchest to parallax-filtered bright field (pBF) and parallax-filtered integrated differential phase contrast (πDPC) and demonstrate their advantages in tomography using T4-bacteriophages, whole cells, and cryo-lift-out lamellae of cellular multilayers. The results show significant improvements over traditional STEM modalities in a realm where conventional wide-field transmission EM imaging methods are not applicable. The commercial availability of such detectors and the relative ease and speed of image reconstruction should make this realm accessible to the broader community in life science EM and beyond.

**Teaser:** Parallax-corrected cryo-STET imaging provides coherent phase and amplitude contrast of thick biological specimens.

## Introduction

Electron microscopy (EM) is the method of choice when tackling ultrastructural studies in cell biology because it reveals all the macromolecular players in the “cellular theatre”. Cryo-EM is an imaging modality whereby the specimen is preserved in a near-native, fully hydrated state by vitrification of the embedding medium. Vitrification provides a snapshot in time and protects the delicate samples from damage due to the expansion of ice crystals upon freezing. Moreover, vitrification avoids the Bragg diffraction that disturbs image interpretation in terms of mass-thickness contrast (*1*, *2*).

In studies of macromolecular structure, the practice of cryo-EM is nearly synonymous with wide-field transmission electron microscopy (TEM) using parallel illumination. A measured defocus generates phase contrast in thin electron-transparent specimens, while a contrast transfer function (CTF) remedies effects of the wave aberration. This practice is coupled with an image processing pipeline known ironically as single particle analysis (SPA) for merging of thousands to millions of individual images for 3D reconstruction, often at near-atomic resolution (*3*). Tomography, on the other hand, generates a 3D reconstruction of a region of interest from a series of projection images acquired at pre-determined tilt angles. Molecular complexes such as ribosomes or viruses may then be reconstructed at high resolution by averaging of small sub-volumes by subtomogram averaging (STA, *4*–*7*), while maintaining the context of their location in the cell (*8*). This context is better maintained in thicker specimens. There is an inherent contradiction, however, because energy loss of the illuminating electrons in passing the specimen couples with chromatic aberration of the magnetic objective lens to create a defocus spread (*9*). An energy filter is used to remove the resulting image haze, but this reduces the image intensity exponentially with a characteristic thickness given by the mean free path for inelastic scattering, about 300 nm for 300 keV acceleration, while still exposing the specimen to damaging radiation. This thickness limitation has motivated the production of thin cryo-lamellae, either by cryo-microtomy (*2*) or, more popularly, by focused ion beam (FIB) milling (*10*–*12*), for example of whole C. elegans L1 larvae (*13*), human forebrain organoids (*14*), or cell-derived matrices (CDMs) (*15*).

An alternative modality for cryo-EM uses a scanned focused probe in transmission, i.e., STEM. Traditionally, the contrast in STEM images originates in the radial distribution of electrons scattered from the sample and recorded pixel by pixel. Rather than recording an image on a camera, one or more detectors integrate the scattering to specific angular ranges while the probe scans across the specimen. The various acronyms for STEM refer to detector configurations, for example, bright field (BF) or annular dark field (ADF) STEM. Such configurations are typically referred to as incoherent contrast methods (*16*), because they record intensities directly. Thereby, phase contrast based on the electron wave coherence is lost. However, STEM largely circumvents the problem of chromatic defocus spread so the energy filter is not required. Thicknesses up to 1 µm or even more are accessible (*17*), and inelastic scattered electrons still provide useful signal. These factors make STEM more dose-efficient than TEM for thick specimens (*17*–*20*). On the other hand, STEM resolution has been, at least to date, inferior to that attainable by phase contrast TEM. Cryo-STET resolution is limited by a number of factors, primarily by the diameter of the focused probe, which is dictated by the convergence angle, α, and additionally by sampling considerations in the raster scan as well as in the tilt intervals. The comparative resolution advantage of cryo-TEM rests on its exploitation of phase contrast, i.e., coherent interference effects in the image formation.

Modern detectors for STEM can be arranged to exploit phase coherence as well as the scattering contrast described above. This is the motivation for ptychography and other methods involving 4D STEM, wherein a 2D diffraction pattern is recorded at every pixel, utilizing a pixilated camera rather than diode area detectors. 4D STEM under cryo-conditions requires a combination of sophisticated equipment that is not yet widely available, and a lengthy data processing pipeline that is still under intensive development (*19*, *21*–*24*). The simplest and most mature phase technique is *differential phase contrast* (DPC), initially proposed in the 1970s by Rose (*25*), and Dekkers and DeLang (*26*), which generates contrast from the difference between scattering in azimuthally opposing directions, utilizing a detector divided into four segments, i.e., a “quadrant detector”, either with or without a central hole (circular or annular). The orthogonal difference signals determine a vector that reflects the gradient in electron phase ∇*ϕ* at the specimen exit face (*27*). These, in turn, reflect gradients in electro-refractive properties, typically at gradients in the electric inner potential of materials. This vector DPC image is in principle a reflection of the local electric field. It may be differentiated to dDPC, which is then a measure of the charge distribution given by the Laplacian of the phase, or integrated in iDPC to provide the phase itself, which is proportional to the local electric potential. Conventional iDPC has recently been applied to biological samples at ambient or cryogenic temperatures (*23*), and sub-nm resolution has been demonstrated for protein structures by SPA (*28*). DPC-compatible quadrant detectors are available commercially and can be installed on modern transmission electron microscopes. Our aim here is to demonstrate the versatility of new data processing methods that exploit these detectors for cryo-STET of biological specimens.

To image thick samples and in tomography, which is the ultimate goal of this study, the beam will never be perfectly focused on the sample. This is especially true for imaging with a high convergence angle which results in a low depth of field. For a defocused beam in STEM, an additional phase due to the incident wave aberration must be considered. With the focus cross-over above or below the sample plane, each point on the specimen is illuminated obliquely from adjacent locations. The relevant angle is an increasing function of the distance from the optic axis; when defocus is the dominant aberration it is simply proportional to the displacement. This oblique illumination creates a parallax that is manifested as a real-space shift, Δl, between images formed independently on each of the DPC quadrants (*29*). The parallax shift on a quadrant detector may be recognized as a simplified case of tilt-corrected bright field (tcBF) imaging introduced recently for 4D STEM (*19*). In both cases, the first step is to de-shift the images computationally as a means of compensating for the defocus. Next, the deshifted images may simply be summed for a coherent parallax bright field result (pBF). Alternatively, iDPC may be computed from the deshifted images to approximate the iDPC or integrated center of mass (iCOM) image that would have been recorded in focus. In our earlier work (*29*), this deshifted iDPC was called iDPC1; for a thin sample it represents the linear phase *ϕ*. The difference between this image and the original iDPC represents the higher order terms extending beyond the weak phase object approximation. These are dominated by the defocus. This difference image therefore represents the depth, determined by parallax, and was originally labeled iDPC2. A further insight, developed here in full, is the role of iDPC differentiation with respect to the parallax correction. Conceptually, this is similar to the classical TEM method of focal pair differences (*30*) and arithmetically it is equivalent to the integration of quadrant signal differences from different heights. Differentiation is achieved by computing the fully parallax corrected iDPC, and then subtracting another iDPC image with the parallax shift deliberately mis-corrected by a small amount. (The choice of such a mis-correction by one pixel, Δ*l* ± 1, corresponds to an axial displacement of one depth of field unit *λ*/*α*^2^, when the pixel sampling matches the lateral resolution *λ*/*α*.) This variational approach proves to be robust in thick samples and is not sensitive to the precise degree of defocus at acquisition. Like iDPC2, it isolates the quadratic contributions to the coherent contrast, also called coherent amplitude contrast. The technique was proposed without elaboration in supplementary material of our first iDPC paper (*29*), where it was called Δ_IS_iDPC (for differential image shift iDPC), and then tested in a 4D-STEM virtual detector simulation of iDPC for cryo-tomography (*23*). In light of the parallax filtering before integration, we introduce the name piDPC, or πDPC.

In the present study, we expand on the theory of πDPC and apply it to data from two commercially available segmented diode detectors. These are the Opal (El Mul Technologies, Inc), which is retrofitted to a Tecnai T20-F (FEI, Inc) S/TEM with a custom scan generator (SavvyScan, *29*), and the Panther detector installed on a modern Talos Arctica microscope (Thermo Fisher Scientific) operating in STEM mode (Fig 1A,B). Data acquisition for the latter required an unconventional scripting, which is described in detail. In addition, we describe the data analysis using a combination of MATLAB and Python scripts. We chose several biological cryo-preserved samples, namely T4-bacteriophages, whole un-thinned cells grown on an EM grid, and focused ion beam (FIB) milled lift-out lamellae of cell-derived matrices (CDM), to showcase a variety of parallax-corrected BF (pBF) and πDPC imaging applications. In all cases, the parallax correction enhances the visibility and sharpness of fine details, without relying on averaging protocols that cannot be applied to unique structures.

**Fig. 1.**
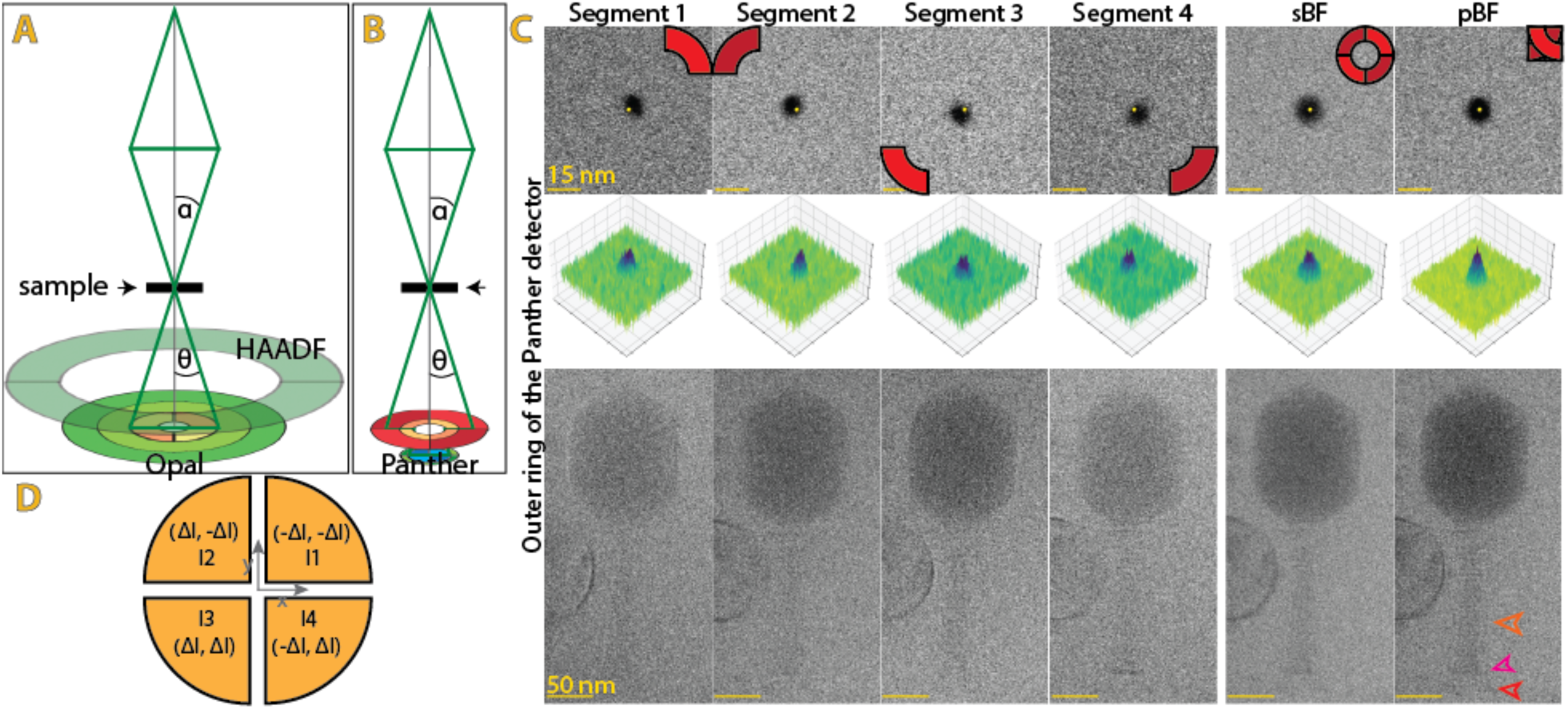
Schematic depiction of the imaging systems and the parallax effect. Schematic of (**A**) the Tecnai T20-F equipped with the Opal and HAADF detectors, and (**B**) the Talos Arctica, equipped with the Panther detector. The green lines depict the schematic BF scattering with the semi-convergence angle, α, and the collection angle, θ. (The top view of the detectors is in FigS1A,B) (**C**) Individual, summed BF and pBF images of a gold bead (yellow dot is in the center of the pBF gold bead) (top row), 3D plot of the intensities (middle row) and a T4 phage (bottom row). The baseplate, the long and short tail fibers are annotated by magenta, orange and red arrowheads, respectively. Additional images of the same gold bead from the three rings of the Panther are in FigS1C. (**D**) Schematic of the four segments depecting the direction of parallax correction. Movie S1 and S2 show the gold bead and T4-phage while cycling through the four segments.

## Results

### Parallax corrected bright field sum (pBF)

2D projection images of gold beads (Fig 1C, top row) highlight the procedure and benefits of parallax correction. A scanned image over the same field of view is produced by each detector element individually. The yellow dot, placed at the center of the gold bead in the pBF (right panel), highlights the fact that these images do not perfectly overlap, even though they were acquired simultaneously in the same scan. When they are simply summed, the gold bead image becomes symmetric around the center but obviously blurred. This simple sum mimics a bright field detector without azimuthal segmentation (simple BF, sBF); the blur is due to the poor spatial coherence recorded by the broad detector, equivalent to illumination by a broad source in conventional TEM according to the theory of reciprocity. A 3D representation of each of the image intensities in the image highlights the relative displacements (center row). The mutual image shifts from the center, Δl, reflect the focus aberration, or, geometrically, the fact that the illuminating rays directed to the particular detector element actually enter the specimen adjacent to the recorded scan location. In magnitude, Δl is proportional to the product of defocus and acquired scattering angle (equivalently, the radial momentum coordinate). The dependence on the collection angle is shown in Fig. S1 by visualization of the individual segments from three rings on the Panther detector. The shift is also similar to a tilted illumination in wide-field conventional TEM (*31*), long recognized as a diagnostic for focus adjustment, and it can be compensated almost trivially by deshifting on the computer. Movie S1 shows an animation flipping through the quadrant images in sequence, where the bead appears to dance around the center point. The improvements of pBF is also shown by focusing on an individual T4-phage and a nearby vesicle in a 2D-projection image. The stalk of the phage in the sBF image appears smeared compared to the images originating from individual segments. This smearing is removed by correcting the parallax shift before summation. This procedure (parallax BF, pBF) reveals the details of the stalk, a long tail fiber, the base, and the collar. A membrane vesicle adjacent to the T4-bacteriophage shows differential shading depending on the angular position of each segment and the directional path of the electrons, similar to directional lightning effects in photography (panels 1-4, from left to right). Simply summing the four segments (sBF) results in a uniform distribution of contrast around the vesicle. However, the membrane is much thicker than in the individual images, whereas the pBF image recovers the sharpness of the membrane of the vesicle. This demonstrates that deshifting indeed improves the resolution and feature sharpness of BF-STEM over use of a single integrating detector. A qualitative justification based on the theorem of reciprocity recalls the theory supporting tcBF (*19*). Parallax correction enables the use of multiple small detector elements, across each of which the wave coherence is better preserved.

### Azimuthal segment imaging on the Opal detector

We have described parallax correction and iDPC calculation on the Opal detector in Seifer et al. (*29*) using a sample of boron nitride nanotubes. Here, we apply the contrast generation methods for advanced cryo-STET reconstruction of mitochondrial membranes in unsectioned tissue culture cells grown directly on the EM grid. A MATLAB script calculates the image shift Δl between the segments, which is proportional to the defocus, and exports sorted tilt series of the sBF, pBF, and πDPC images (and additionally the iDPC, iDPC1, and iDPC2). For a detailed description of πDPC, see the Materials and Methods section. Briefly, it is the integration of DPC computed from the parallax correction suited to two nearby planes, essentially a derivative along the defocus axis, or physically the direction of depth. The tomographic tilt series is then aligned using standard software, e.g., IMOD or AreTomo, to produce a transformation file that approximates rigid body rotation around a unique axis, identically to conventional TEM tomography. For this purpose, we prefer to use the pBF; as the simplest approach to correct for defocus blur, it is typically the most robust. The transformations are applied immediately via a script to the other contrast mechanisms, which reconstructs the tomograms using standard back-projection, followed by 3D deconvolution (*23*, *32*, *33*). All the STEM contrast modes written out by the script are shown in Fig. 2 (sBF, pBF, and πDPC) and Fig. S2 (iDPC, iDPC1, and iDPC2).

**Fig. 2.**
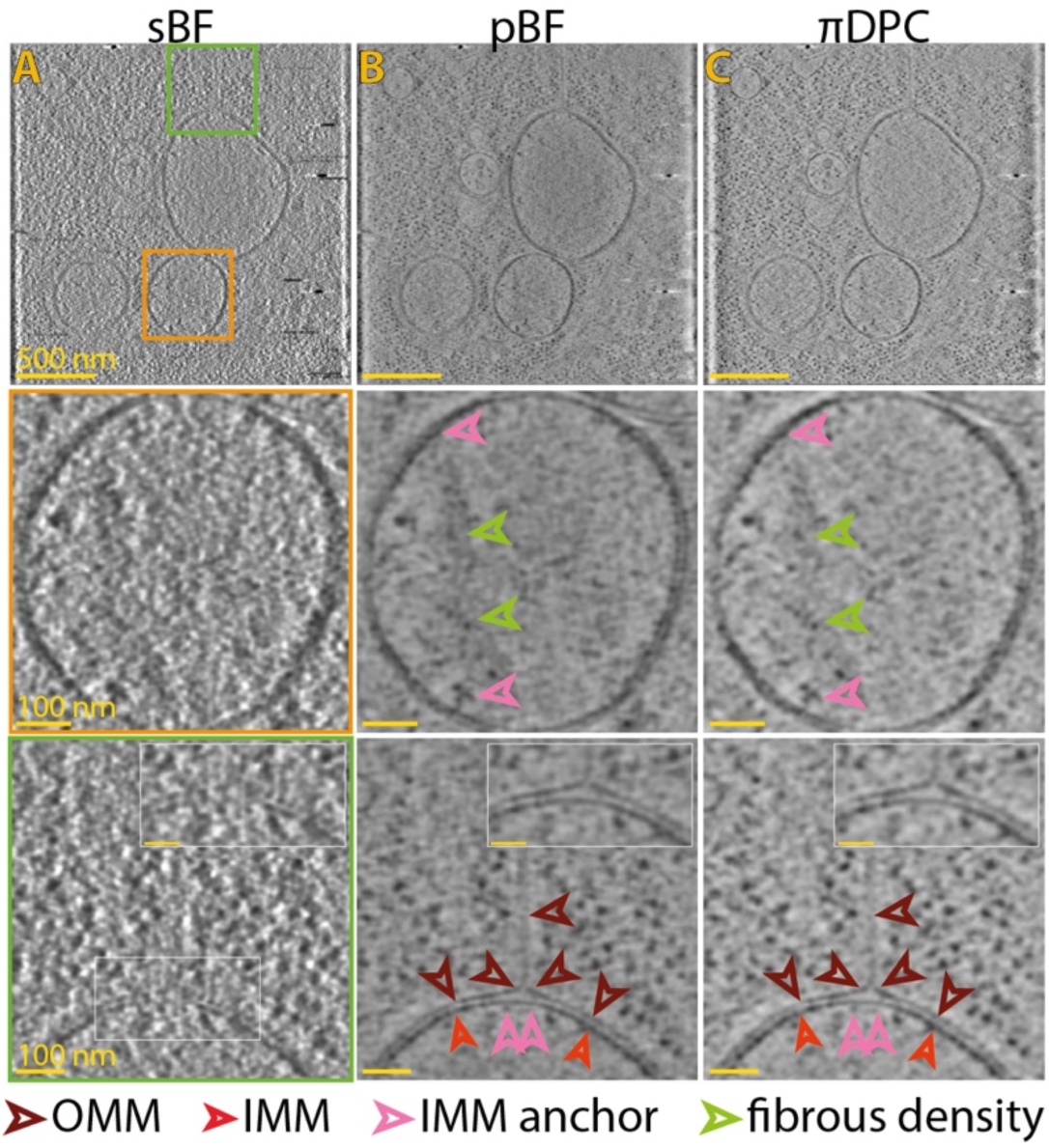
Comparison of the sBF, pBF and πDPC. Individual virtual slices of a tomogram recorded on the Opal detector, reconstructed in the (**A**) sBF, (**B**) pBF, and (**C**) πDPC mode, showing three mitochondrial fragments (top row). The orange and green boxes highlight areas magnified in the middle and lower rows, respectively. The middle row zooms into the one of these mitochondrial fragments. The lower row zooms into the formation of a mitochondrial nanotunnel pinching off of the outer mitochondrial membrane. Scalebars are 500 nm, 100 nm, and 100 nm, in the top, middle, and lower rows, respectively.

The different contrast mechanisms from the Opal are highlighted in Fig. 2 and Fig. S2. We show some mitochondrial fragments recorded within an intact cell with a thickness around 400 nm. The sBF shows the mitochondrial membrane, with the outer (OMM) and inner mitochondrial membrane (IMM) being visible. However, details inside the mitochondria (second row, orange box) and even membrane continuity (third row, green box) are difficult to see. pBF and πDPC reveal details in the matrix of the mitochondria (middle row). Due to the irregular shape and the anchoring at the IMM, we suspect these are parts of a mitochondrial DNA molecule (mtDNA). Additionally, a mitochondrial nanotunnel is revealed (bottom row). Interestingly, the contrast coming from the uncorrected iDPC is intense but fuzzy (Fig. S2). When separating the parallax-corrected iDPC1 from the defocus-dependent iDPC2, it becomes evident that iDPC2 allows restoration of the high-quality image, while the fuzzy contrast comes from iDPC1. This reproduces our earlier observations on an room-temperature sample on the Opal detector (*29*), and 4D-cryo-STET on a DECTRIS ARINA camera (*23*), that iDPC2 is more useful than iDPC1 for phase contrast in continuous-volume samples such as cells. It is not surprising that πDPC and iDPC2 produce similar contrast because they differ in the magnitude of the discrete differential step (1 pixel shift or the difference between recorded defocus and the shift correction to ideal focus). It is somewhat surprising, nonetheless, that the pBF and πDPC images are so similar after tomographic reconstruction.

### Azimuthal segment imaging on the Panther detector

We applied the cryo-iDPC workflow next to the Panther detector. This detector enables data collection using twelve segments simultaneously: two annular rings and one central disk, all of which are divided to four segments azimuthally. At the time of this writing, the manufacturer’s tomography software does not support export of data from all the individual segments. Therefore we saved the data from each segment via a custom protocol involving SerialEM (*34*) to operate the microscope, AutoscriptTEM (ThermoFisher) to collect the data, and the Python mrcfile library (*35*) to write out the data array in standard image file format. We adapted the parallax correction & iDPC MATLAB script to calculate the different contrast mechanisms for each ring of the detector individually. For proof of principle, we targeted mitochondria from U2-OS fibroblast cells. The 0-degree view of the tilt series from the outer ring exemplifies the deshifting effect. While the outline of the mitochondria is visible in the sBF (Fig. S3A), the two mitochondrial membranes only become visible after parallax correction (Fig. S3B). Finally, we merged the data from the three rings, used the pBF image to align the tilt series, and applied the transformations to the individual rings and the other contrast mechanisms.

The 3D volume shows several large mitochondria in an area more than 700 nm thick (Fig. 3). Close to a fission site, is a mito-ER contact site with several mitochondria-ER tethers (Fig. 3, orange box). These tethers are different in shape, some with one adaptor, while others with two adaptors. When comparing the pBF to the πDPC in the combined image (Fig 3), or just the outer ring (Fig. S4), no major differences in details and the resulting contrast are visible. Many of these features are seen the in the weighted back projection tomogram of the pBF (Fig S5A) and πDPC (Fig S5B), however, deconvolution increases the contrast significantly, as described previously (*32*, *33*). The πDPC may also be computed using individual rings on the Panther (Fig S5). At the low dose rates used for tomography, not much signal is collected by the innermost detector for such a thick sample. Contrast in the middle ring is very strong. However, the outer ring seems to show the finest features. This is entirely in accord with expectations that the highest spatial frequencies are reflected at the highest scattering angles.

**Fig. 3.**
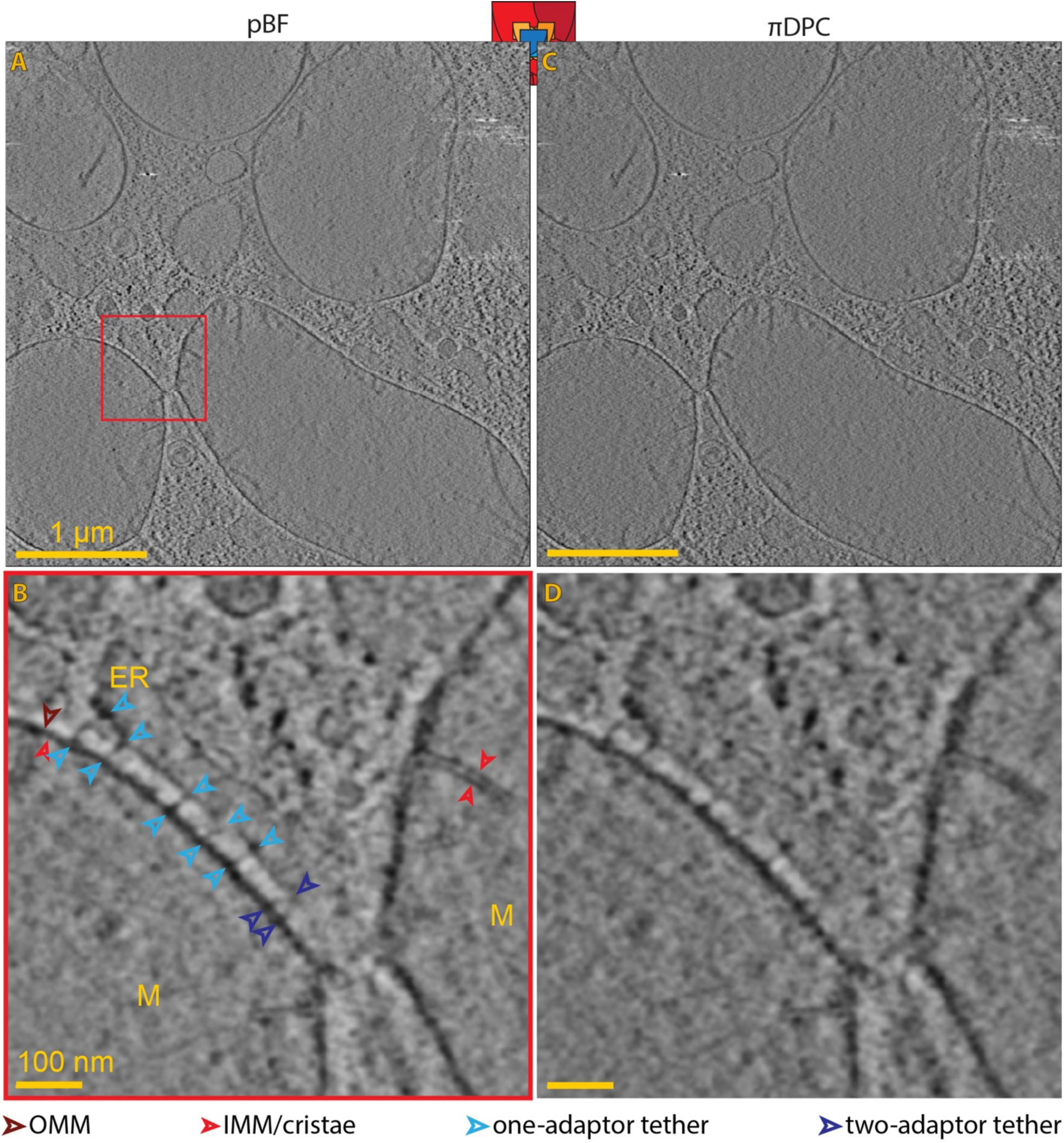
Comparison of pBF and πDPC from the Panther detector. A single slice of a 700 nm thick tomogram from the (**A,B**) pBF and (**C,D**) πDPC showing several mitochondria from U2oS cells. Zooming into a (**B,D**) fission side (red box), highlights a mitochondria-ER contact site, showing several tethering proteins. Scale bars are 1 µm and 100 nm in the top and bottom panel, respectively.

When comparing the different iDPC contrast modes (Fig. S7), it becomes obvious that pBF and πDPC (Fig. 3) provide a more interpretable contrast than iDPC (Fig S7A), iDPC1 (Fig S7B), or iDPC2 (Fig S7C) for the thicker cellular samples.

### Cryo-STET of cryo-lift-out lamellae

A major recent development in specimen preparation for cryo-tomography is that of FIB-milled lift-out lamellae. By extracting a small portion of the sample and laying it on a grid, the method may be applied to CDMs (*15*), tissues (*14*), or even whole organisms (*13*). Cryo-STET should then enable observation of thicker sections than conventional EFTEM tomography. In order to test this expectation, we employed CDMs, in which human telomerase immortalized foreskin fibroblasts (TIFFs) form cell multilayers and produce their own extracellular matrix (ECM) environment (*36*). The resulting ECM consists of fibrous proteins (e.g. collagen) and proteoglycans (*15*). Lamellae were prepared as previously reported (*15*), but milled to an intended thickness of around 600-900 nm. The lamellae were taken from the same sample as in the original publication.

First, we imaged the CDM-lamellae using the Tecnai/Opal system and processed the data to generate the pBF and πDPC contrasts (Fig. 4A,B, and movies S1 and S2, respectively). The field of view was approximately 2 µm in x and y, with thickness 400 nm. In the tomogram, we see two cells in cross-section separated by the extracellular space filled with protein fibers, presumably collagen. The cell on the left shows a nucleus, identified by the double bilayer membrane of the nuclear envelope. A gap of appropriate size in the membrane suggests a nuclear pore complex (NPC) (Fig. 4B,E, cyan box). By way of confirmation, we manually placed a model of the human NPC generated by subtomogram averaging onto the membrane, and note nearby densities reminiscent of the cytoplasmic ring and nuclear basket ((EMDB-ID: 3103, *37*) Fig 4E). Within the nucleus we observe a thin fibrous density punctuated by beads of about 10 nm diameter, as would be expected for decondensed chromatin (Fig. 4C,F, orange box).

**Fig. 4.**
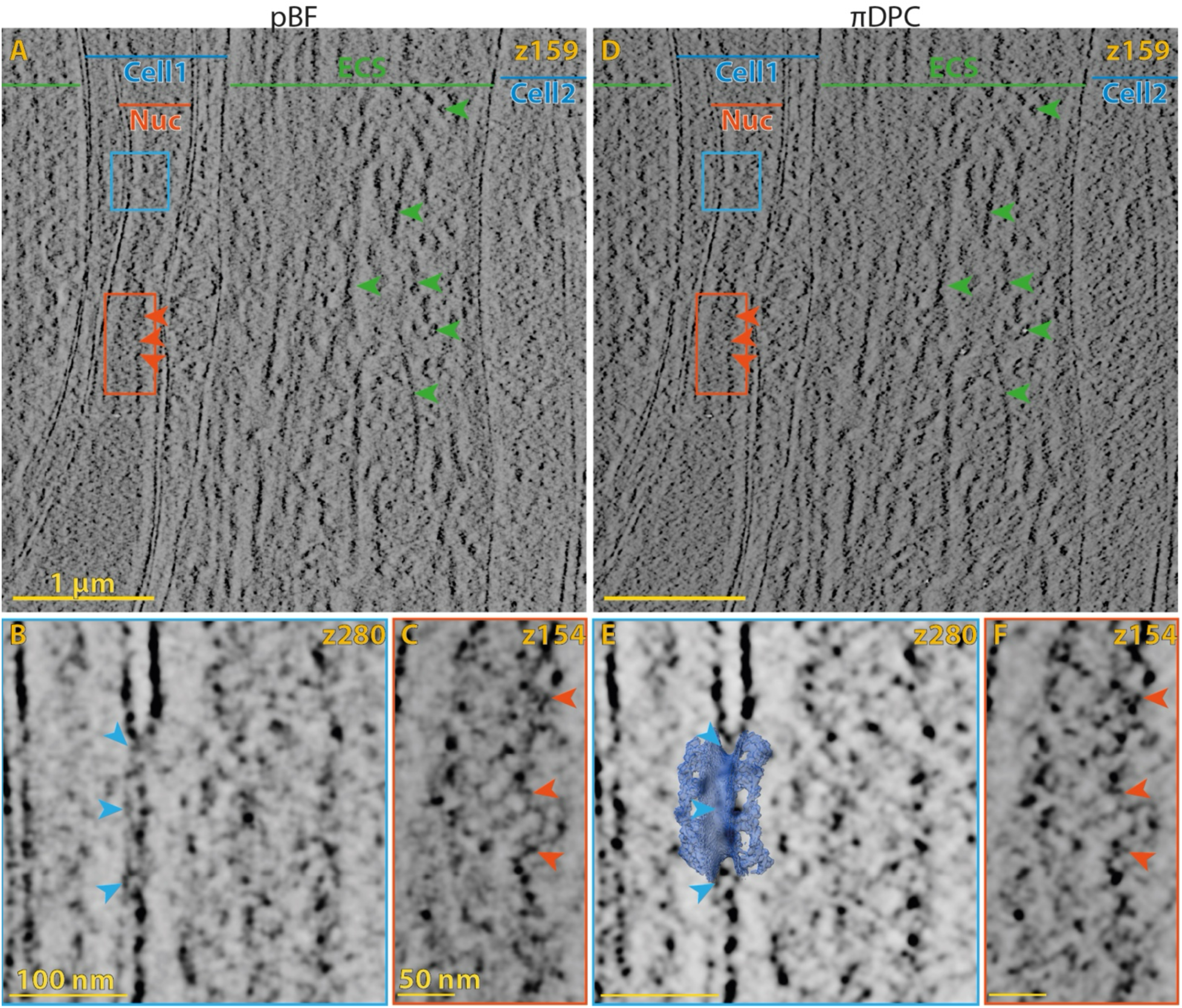
Comparison of pBF and πDPC imaging on lift-out lamellae from CDMs on the Opal detector. A single slice of a 400 nm thick tomogram from the (**A-C**) pBF and (**D-F**) πDPC showing a lift-out lamellae. The data shows two cells separated by the ECM, which is filled with fibrous densities (green arrowheads). Blue and orange boxes highlight a potential position of a (**B,E**) nuclear pore complex (blue arrowheads, overlayed is the EMDB-3103 (blue) (*37*)) and a (**C, F**) chromatin strand (orange arrowheads), respectively. Scale bars are 1 µm and 100nm/50 nm for the overview image and the zoom-ins, respectively.

We then repeated the demonstration with a second sample using the Arctica/Panther system. The specimen in this area of the lamella was 700 nm thick, and we recorded tilt series with a field of view of 4 µm in x and y. Virtual sections of πDPC contrast show several layers of cells interspersed with ECM (Fig. 5, movie S3, Fig. 6, movie S4). Segmentation gives an impression of the 3D-nature of the data. In some instances, we see the cell membrane and collagen fibers in close proximity, potentially indicative of an interaction between these two entities (e.g. Fig 5C). In the first tomogram, we find mitochondria in contact with the rough ER (Fig 5D-F). In the second, we find an endocytic site in cell 2 (Fig. 6C). Cell 3 contains many intracellular granules (pink arrowheads).

**Fig. 5.**
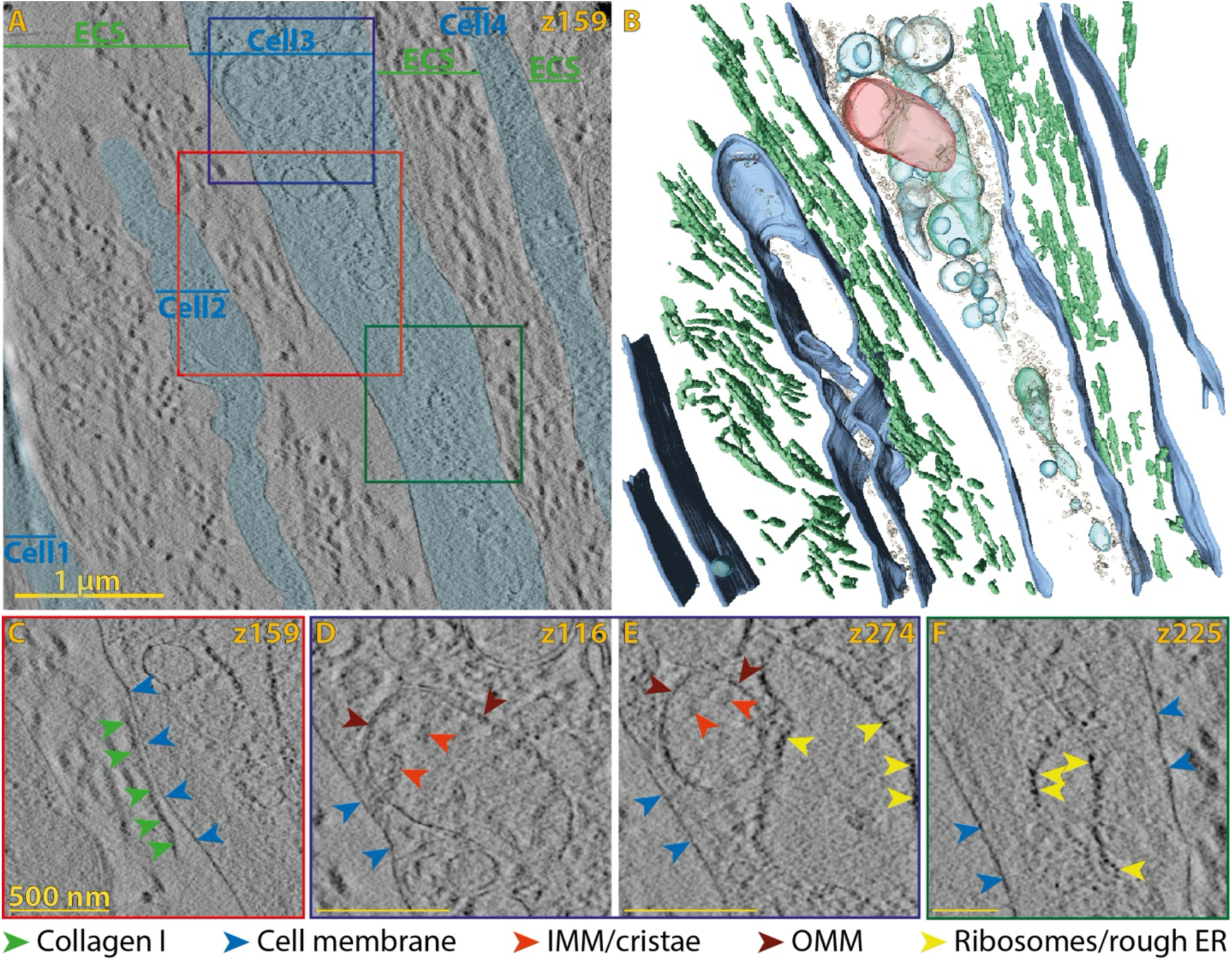
πDPC imaging of the CDMs from the Panther detector. (**A**) shows the 159 slice in the tomogram (with the cellular parts shaded in light blue) and (**B**) the 3D segmentation. Blue, green, cyan, red and grey show the cell membrane, collagen fibers, intracellular vesicles (including ER), mitochondria and proteinous structure, respectively. (**C-F**) show selected areas with the z-slice indicated. Scale bars are 1 µm and 500 nm for the overview and the zoom-ins, respectively.

**Fig. 6.**
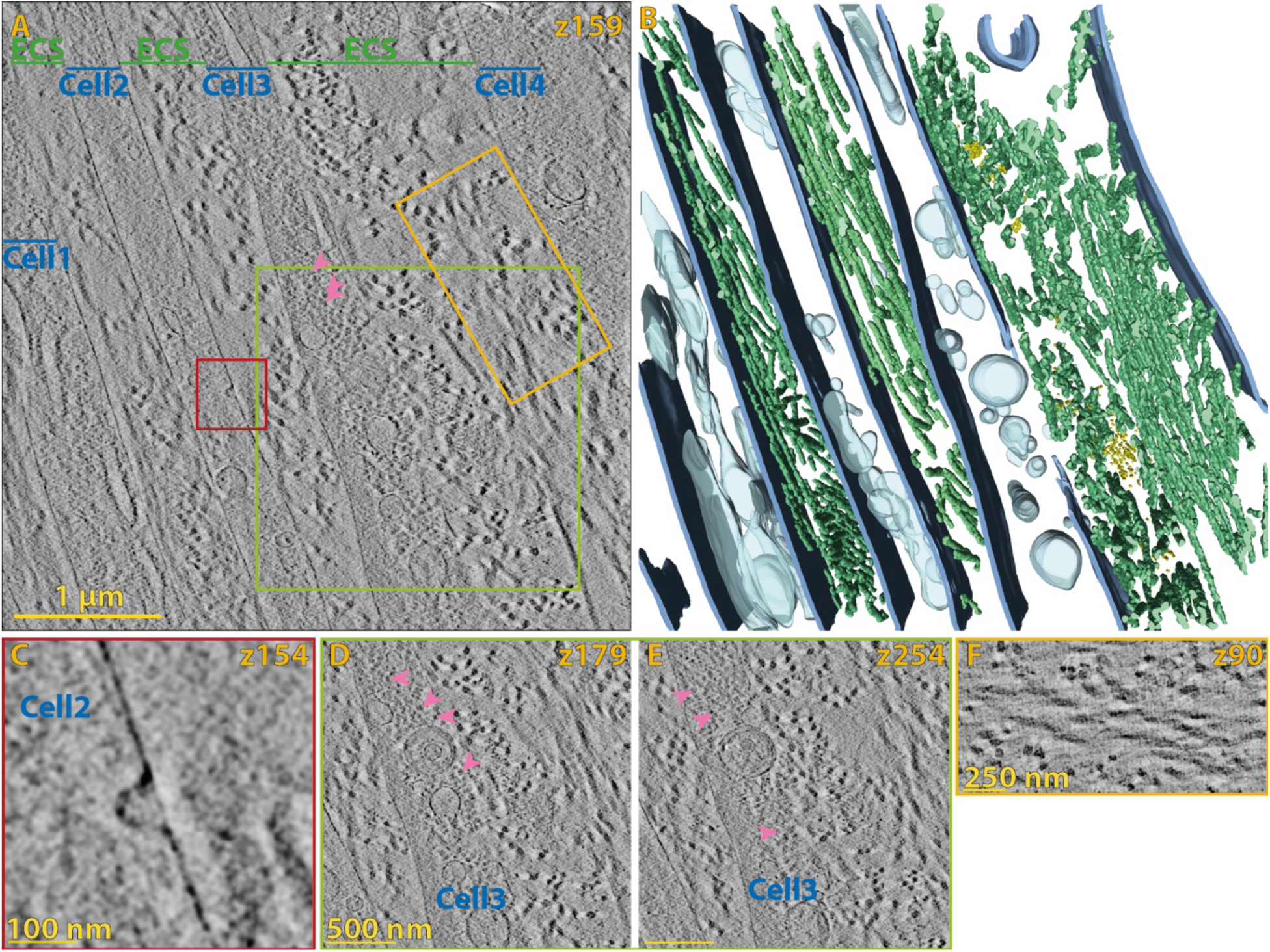
πDPC imaging of the CDMs from the Panther detector. (**A**) shows slice 159 in the tomogram and (**B**) the 3D segmentation. Blue, green, yellow, and cyan show the cell membrane, collagen fibers, smaller ECM fibers, and intracellular vesicles (including ER) respectively. (**C-F**) show selected areas with the z-slice indicated. Pink arrowheads show intracellular granules. Scale bars are 1 µm in (A), 100 nm in (C), 500 nm in (D+E) and 250 nm in (F). Movies S

The extended field-of-view afforded by Cryo-STET allowed us to analyze the orientation of the fibers in the ECM area (Fig 7). The fibers were divided into four groups: Group 1 (ECM area between cell 1 and cell 2, 85 fibers), Group 2 (ECM area between Cell 2 and Cell 3, 56 fibers), Group 3 (ECM area to the right of Cell 3 contains 134 fibers), and Group 4 (ECM area to the left of Cell 4, 61 fibers). We then extracted the angles they form with respect to the z-axis (Φ-angles) and present them as violin plots. In all groups, we find individual fibers covering the entire range from 0-90°. However, the majority of fibers, as highlighted by the width of the plot, are arranged differently for each group. The predominant orientations are seen as widening of the plots, respectively for groups 1-4 at around 50°-70°, 20°-40°, 5°-20°, and 50°-70°. The difference in the fiber orientation between groups 3 and 4, even though they share the same space in the ECM, is highlighted in the inverted contrast stereo view (Fig 7C).

**Fig. 7.**
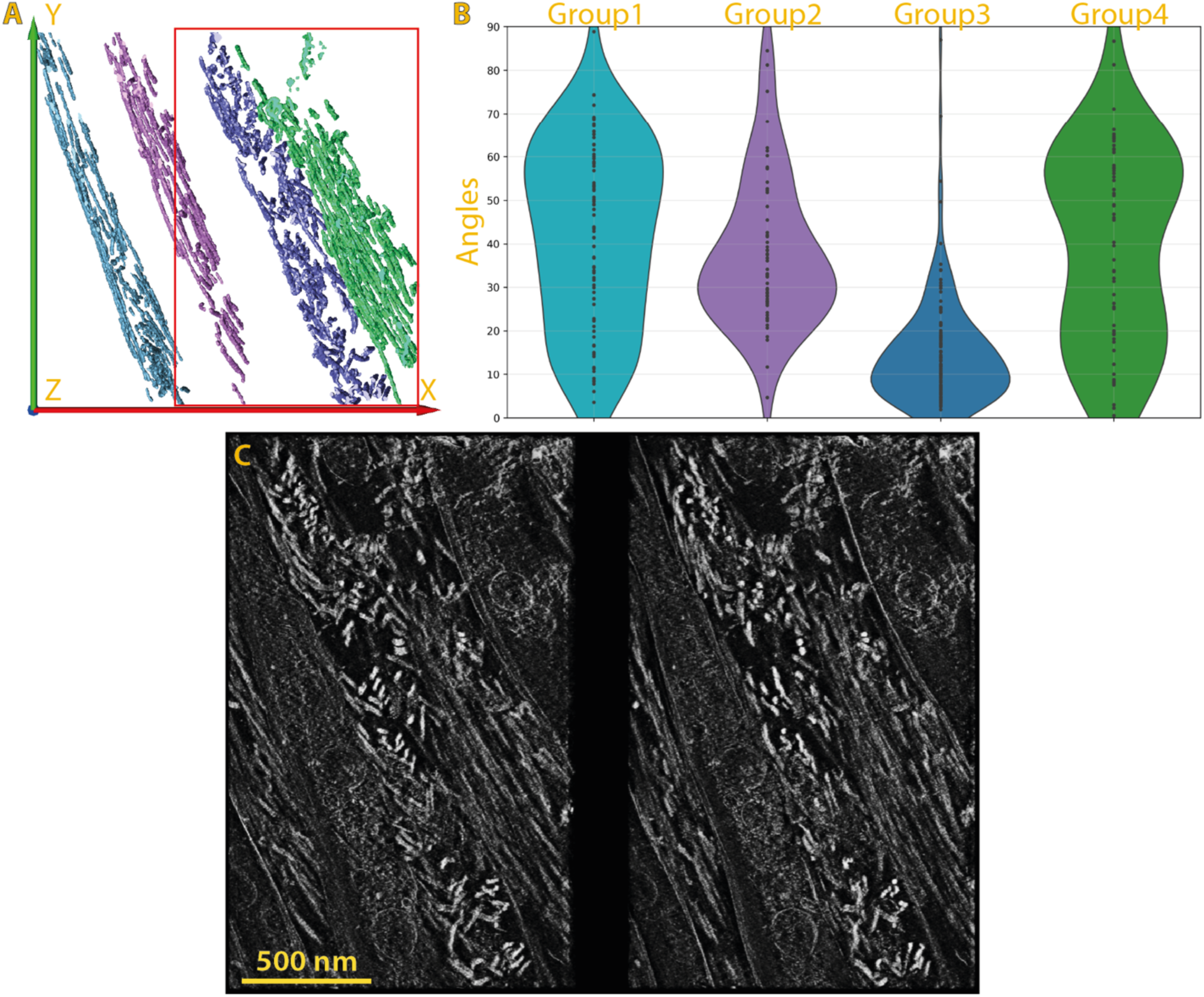
Orientation statistics of the collagen fibers from Fig 6. (**A**) Overview of the four groups. (Red box shows view in (C)). (**B**) Violin plot showing the distribution of fiber Φ-angles relative to the z-axis, with the violin’s width representing kernel density and black points indicating individual measurements. (**C**) An inverted contrast walleye stereo view of the neighboring ECM area of group 3 and 4. The view convers slices 20-320 of 347 slices. The scale bar is 500 nm.

## Discussion

Cryo-electron tomography generates a 3D reconstruction and visualization based on a series of 2D projection images. It is widely applied to biological specimens, where vitrification provides faithful preservation of the delicate organic matter. Cryo-tomography studies then follow two branches: one is the approach of *in situ* structural biology, aiming for high resolution molecular structure in the spirit of SPA but in the native cellular context, and the second puts the primary emphasis on the cellular structure itself. In the latter direction, much effort has been dedicated to expanding the reconstructed field of view by fast acquisition of large area montages (*38*–*41*), often with an eye to finding rare targets for STA. In TEM-based cryo-ET, this often results in a volume of 0.5 µm^3^, mainly limited in the attainable thickness, while STEM-based cryo-ET, as shown here, the volume reaches 13 µm^3^. While montaging can expand the lateral field of view, the specimen thickness remains limited in wide-field TEM by energy loss and chromatic aberration. Conventional STEM configurations largely circumvent that limitation, but lacking the benefits of phase contrast they were clearly inferior in terms of resolution. A SPA study of zinc decorated ferritin showed only 2 nm resolution for the protein structure, but better than 3Å uncertainty in the location of the metal atoms (*42*), suggesting that much higher resolution should be attainable. More recently, sub-nanometer resolution was indeed demonstrated by SPA for protein structures using iDPC (*28*) or STEM-based ptychography (*24*).

Here we have presented the optimization of parallax correction for iDPC cryo-tomography, together with examples that were out of reach by conventional cryo-ET. While iDPC has been widely adopted in material science, it is disturbingly sensitive to the precise focus conditions for high resolution (*43*). The same sensitivity and loss of contrast with defocus may be exploited, on the other hand, for optical sectioning without tomography (*44*). To the extent that defocus represents a simple wave aberration in the illumination, it results in a translational shift between the off-axis elements in a segmented detector. Compensating the shift computationally has an effect similar to refocusing. After deshifting, the sum provides an improved image, the pBF. For isolated weak phase objects, the iDPC computed on the deshifted data produces a phase image with extended depth of field, while the remainder yields contrast according to depth, as could be seen with a nest of isolated inorganic nanotubes (*29*). An improvement on this difference is the differential with respect to defocus, which we call here πDPC. We find it more reliable than conventional iDPC in that does not depend so sensitively on the initial acquisition conditions. More technically, the deshifted iDPC and its differential with respect to image shift have the effect of separating the linear (coherent phase: imaginary and sinusoidal) and quadratic (coherent amplitude: real and attenuating) contributions to the contrast (*29*). A thick, continuous specimen, such as a part or section of a biological cell, is normally not a proper weak phase object, and we find here that the best interpretable signal is that of the coherent amplitude contrast recorded in the bright field.

Judged visually, the tomographic sections reconstructed by pBF and πDPC methods appear remarkably similar. We can surmise that this is due to the predominance of the coherent amplitude contrast for thick samples. Considering the theorem of reciprocity, pBF is similar to an axial STEM image with intensities added after accounting for the relative image shifts. Forward scattering to higher angles in the coherent beam electron diffraction (CBED) paradigm reflects a smaller scale and finer detail. On the Panther detector, for example, each concentric disk or ring contributes to a different range of spatial frequencies in the image representation (Supplementary Fig S5). iDPC, on the other hand, works from the intensity difference recorded from opposing detector segments. In analogy with Bragg scattering, one may expect that the scattering to opposite directions in the diffraction plane should carry opposite phase so that the sum and difference reflect Friedel and anti-Friedel components. This distinction is likely limited to the weak linear phase contrast that is poorly reconstructed in the tomograms (iDPC1 in Supplementary Figs S2 and S4). For such thick specimens the contrast is dominated by the quadratic term described in the theory section, which is similar in both cases. An important distinction between pBF and πDPC is that the latter suppresses the influence of incoherent scattering to the high angle dark field regime, whereas pBF retains that signal.

The parallax correction employed here with a quadrant detector bears a strong resemblance to the tilt correction proposed for 4D STEM using a pixelated detector (*19*). That approach, tcBF, implements a correction of the defocus by deshifting according to the vector position in the diffraction plane. The pBF presented here is conceptually similar to tcBF with four “pixels”. tcBF recovers a sinusoidal contrast transfer function (CTF) similar to that of conventional wide-field TEM. A further extension of the tilt correction has been proposed to tcDPC (*45*), which appears to be similar to the parallax-corrected iDPC1 described in (*29*). The πDPC described here is a variation in the tilt correction, as a differential with the parameter of defocus. In another work, we have shown that for sufficiently large defocus the tcBF is equivalent to a montage of overlapping shadow images (*46*). Both methods are able to reconstruct a real-space image at higher resolution than the originally recorded pixel sampling, based on interpolation from the multiple diffraction plane images. The shadow montage method was applied also to tomography. With only four “pixels”, the quadrant detector data is not suitable for such up-sampling. On the other hand, the quadrant detector read-out is much faster than that of the 4D STEM camera and the datasets are much smaller, facilitating the processing.

To highlight the new cryo-STET method we focused on thick specimens and features that are not immediately amenable to subtomogram averaging. These included mitochondrial membrane structures and a fibrous inclusion that we suspect may be the elusive mitochondrial DNA. Further, we were able to visualize the organization of ECM protein fibers that spanned the spaces between cells in CDMs. Mass spectrometry revealed a that the fibrous ECM is composed of primarily of collagens I and VI, and fibronectin (*15*). Based on size measurement made by conventional cryo-ET, the fibers appeared to be predominantly collagen I, with some smaller filaments present as well as spherical granules and an amorphous matrix nearby fibers (*15*). The present data are consistent with that determination; relatively thick and thin fibers are highlighted in Fig 6B. The extended specimen depth accessed by cryo-STET allows following ECM fibers over longer distances, hence enabling a more thorough determination of the molecular arrangement of cells and ECM components, such as out of plane tilt angles (Φ-angles) with respect to cells.

In summary, we have shown that parallax processing is a powerful enhancement to cryo-STET. This builds on prior work incorporating BF-STEM, 3D deconvolution and dual-axis tomography for visualization of whole mitochondria (*20*, *33*), chromatin at the edge of a nucleus (*32*), or whole malaria parasites after infection of red blood cells (*47*). The pBF and πDPC methods can both be implemented from the same dataset, allowing for optimization to the case at hand, and they relax the requirement in iDPC for precise focus control. They produce similar contrast in thick (400-800 nm) biological samples. Differences between them would likely appear for thin samples and at higher resolution than what is achievable here. Parallel developments using 4D STEM pixelated detectors offer a technological alternative (*19*, *23*, *24*, *46*). Comparatively, the quadrant detectors are simpler to implement and the hardware is more widely available. This study demonstrates the parallax implementation with widely available software controlling the microscope, as well as a partially automated data analysis. The workflow employed here to visualize CDMs, namely prolonged growth of fibroblast cells, induction of ECM secretion, high-pressure freezing, cryo-lift-out lamellae preparation and cryo-STET imaging, could equally be applied to other higher-order samples such as organoids, or whole organisms as well and opens many possibilities to visualize biological processes in 3D.

## Materials and Methods

### Sample preparation for EM

#### T4-Bacteriophages

Cryo-preserved T4-bacteriohages were isolated as described previously described (*23*). In short, E. coli were cultured to propagate the bacteriophages at a multiplicity of infection of 1. After 5h of incubation, the T4 phages were enriched by filtration and differential centrifugation. As a first step, the remnants of the bacteria were pelleted for 30 min at 5000g. The supernatant was then filtered twice using a 0.46 µm and a 0.22 µm pore filter. The T4-Phages were then pelleted at 35.000g for 40 min at 4℃. The pellet was resuspended in 50 mM Tris pH 8 with 1 mM MgCl_2_ and filtered through a 0.22 µm filter.

The T4-Phages were vitrified by plunge freezing using a Leica EM-GP plunger (Leica Micosystems, Vienna). The chamber was set to 90% humidity and 23℃. A Quantifoil R1.2/1.3, with a 2 nm continuous carbon layer on a 200 mesh copper grid, was glow discharged for 60 sec (PELCO easiGlow). 3 µL of T4 phage samples were added to the carbon side, and 1.5 µL of 15 nm home-made gold fiducials were added from the back side. The grids were blotted from the back for 3.5 sec and stored in liquid nitrogen until usage.

#### *U2OS* cells

U2OS cells were maintained in DMEM (Gibco) + FCS (10% final concentration), + Pen/Strep (1%), + L-Glu, + 1mM Sodium Pyruvate. On the day before plunge freezing, the cells were seeded on Gold Embra Quantifoil R2/4 EM grids. The cells were frozen by plunging into liquid ethane using a Leica EM-GP plunger (Leica Microsystems, Vienna, *48*). Settings during plunging included 37 ℃ at humidity 90%, with blotting from back-side for 5 sec. 3 μL of media samples was added to the carbon side of the EM grid, and 2 μL of homemade 15 nm gold fiducials was added to the backside before plunging.

#### CDM preparation

The preparation of the CDMs lift-out lamellae was described in detail previously (*15*). In short, telomerase immortalized foreskin fibroblasts were grown for three weeks on cryo-EM grids. To induce ECM production ascorbic acid was added. The samples were high-pressure frozen with a BAL-TEC HPM010 and stored in liquid nitrogen until lamellae preparation.

Cryo-lift-out lamellae were generated using a second-generation Aquilos (Aquilos II) instrument (Thermo Fisher Scientific). The instrument was operated using the xT user interface and the MAPS 3.14 software (TFS). The FIB was operated at 30 kV, and the milling progress was monitored using the SEM beam at 25 pA and 5 kV. A 45° pretilt shuttle was used for all cryo-FIB milling steps described below.

Prior to sample loading, a 100/400 rectangular mesh grid (#G1040-Cu; Science Services) was clipped into an AutoGrid as receiver grid. For clipping, the long side of the rectangle mesh was placed perpendicular to the milling window to allow for easy lift-out attachment and low-angle sample thinning.

After loading the sample and receiver grid, overview maps of the high-pressure frozen specimens were acquired and correlated with the images acquired on a Leica EM Cryo CLEM microscope to identify regions of interest. The milling slot on the FIBSEM AutoGrid was used to improve correlation by using its rim, visible in both reflected light microscopy and SEM, as a landmark.

The specimen was cleaned from contaminations such as debris and ice in eucentric position at 20° stage tilt with the FIB operated at 1 nA. Directing the beam to areas with contamination facilitates its removal to create a smooth, clean surface for subsequent coatings. The specimen was sputter-coated with platinum for 30 s at 30 mA and 10 Pa with the built-in sputter coater, followed by a GIS deposition of 1.5–2 µm metalorganic platinum and another sputter-coating (30s, 30mA, 10 Pa). Tile-scan overview images were taken after each step using the MAPS software.

Lift-out sites were identified by CLEM and set to eucentric position. Steps performed for the lift-out FIB milling are explained in greater detailed in (*15*): Trenches for the lift-out procedure were cut at a stage tilt of 7° and a relative stage rotation of 180° to position the FIB perpendicular to the sample. The trenches in front, behind, and to the side of the lift-out were milled in cross-section (CS) patterns with 5 nA. The front and back of the lift-out were polished smooth by milling with cross-section-cleaning (CSC) patterns fitted to the width of the lift-out with 1 nA. Undercuts were performed at 28° stage tilt, at the default stage rotation, with 1 nA with rectangle milling patterns. The micromanipulator needle was then attached by redeposition milling, using a CS pattern with a Multi-Pass setting of 1 at 0.5 nA. The remaining anchor to the bulk sample was removed at 0.5 nA with a rectangle pattern placed at the anchor position. The lift-out was then lifted out of the bulk sample and transferred to the receiver grid. The grid bars at the attachment site were pre-milled at default stage rotation, 22° stage tilt, with a CS pattern and a current of 15 nA. The patterns were set up to produce an angled recess matching the width of the lift-out and a z-depth of ∼15µm. This ensured a better attachment of the lift-out as well as a pre-tilt close to the desired final milling angle.

The lift-out was then attached to this recess at eucentric height at a relative stage rotation of 180° and a stage tilt of 22°. The lift-out block was maneuvered into the pre-milled slot before attachment by redeposition milling, using CS patterns at 0.5 nA with a Multi-Pass setting of 1, a z-depth of 3 µm, and an area of ∼2/1.5 µm (x/y). Following visual confirmation of the attachment, the needle was pulled off gently to the side, so it could be directly reused for the next lift-out without any necessary cleaning steps.

Each lift-out was then thinned with rectangle patterns in several steps, reducing lamella width and milling current in each step, resulting in a symmetric stair-like anchor. Final lamella width was approximately 20 µm. Lamellae were thinned down to 3 µm thickness with 1 nA, then to 1.5 µm thickness with 0.5 nA, and to 900 nm using 0.1 nA. Here, the stage was tilted to ±1° and the lamella overhangs above and below were milled with 0.1 nA to facilitate a parallel shape of the lamella rather than a wedge-like one. The lamellae were then milled down to a target thickness of 600-900 nm with 50 pA.

All samples were stored in Autogrid boxes in liquid nitrogen until shipment to the Weizmann Institute, Israel, for cryo-STET imaging.

### Dose calculations

The electron dose in biological cryo-tomography, usually around 100*e^-^* Å^−2^ to 120*e^-^* Å^−2^, limits the amount of electrons on the detector. The dose per beam position in STEM is calculated as the total number of electrons passing through the sample.

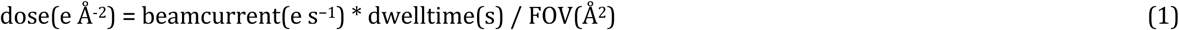

where the beamcurrent is read from the main Screen in nA which can be converted using the dose rate constant 1pA = 6.24×10^6^*e^-^* s^−1^ = 6.24*e^-^* µs^−1^. To calculate the dose for a tilt, the equation changes to

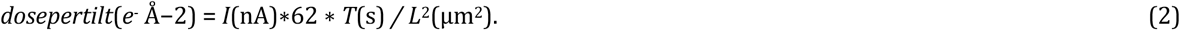

To calculate the dose for the whole tilt series we include the number of tilts, *N*, and change the equation to

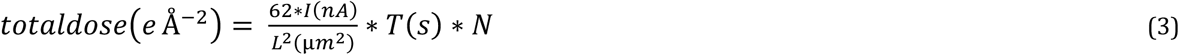

Here *I* is the screen current in nA, *L* is the FOV in µm, and *T* the frame time per FOV in s.

### STEM imaging

A Tecnai F20 microscope (FEI, Inc) was operated in STEM mode at 200 kV as reported previously (*23*, *29*). The microscope is equipped with an HAADF (Fischione model 3500) and Opal detector (El Mul technologies, Israel). The Opal has three rings, with the inner ring being split into four segments. A 30 µm condenser aperture was used to achieve a convergence angle of 0.8 mrad. The spot size and dwell time settings were chosen as to achieve 180 e^-^/A^2^ over the whole tilt series. The specimen was maintained in a Gatan 626 single-tilt cryo-holder (Gatan, USA).

SerialEM (*34*) is used to operate the microscope, and the HAADF channel was used to control tilt series acquisition. Tilt series were acquired in dose-symmetric scheme (*49*) from 60° to -60° in 2° steps. Data collection on the Opal has been described in more detail previously (*23*, *29*). We chose a nominal magnification of 40000, for 2048 pixels, this resulted in a step size of 11.9 Å while recording a FOV of 2.44 µm.

A Talos Arctica microscope (Thermo Fisher Scientific, Inc.) was operated in STEM mode at 200kV as reported previously (*50*). Insertion of a 20µm C2 aperture in nanoprobe mode results in a semi-convergence angle *α* of 2.4 mrad, and a 50µm C2 aperture in microprobe mode results in a semi-convergence angle *α* of 1.2 mrad. Data were acquired on a Panther segmented detector (TFS). The Panther detector has four different rings with four segments each; however, only three rings are usable simultaneously (*28*). We chose a camera length (CL) as to maximize the bright-field illumination signal under vacuum on the outer ring (Fig. S1, B, C). In nanoprobe mode, we chose 3.6 m, and in microprobe mode 5.7 m. To image the T4-phages, nanoprobe mode was used, using a magnification of 71000, resulting in a step size of 2.154 Å, yielding a FOV of 0.8 µm. To image the mammalian cells and the CDMs, the microscope was set to microprobe, and the magnification was set to 28500, for 4096 pixels, this resulted in a step size of 10.81 Å while recording a FOV of roughly 4.4 µm.

The microscope was operated using SerialEM (*34*). The tilt series were acquired using the dose-symmetric scheme from -60° to 60° in steps of 2°. To acquire the 12 segments of the Panther detector individually, the AutoScript TEM Server (TFS), running the Microscope computer, was accessed. For that purpose, a Python script from the AutoScript templates was adapted to record the data, and the mrcfile Python library (*35*) was used to store the data coming from the individual segments in the data folder for each tilt series. The script is found on GitHub, and on the Nexperion script repository. How to use the script is described in more details in the Supplementary information and on GitHub.

### Parallax-iDPC Theory

In theory, the electron is a de-Broglie wave. On transmission through the sample, the wavefront undergoes a position-dependent phase delay proportional to the projected electrostatic potential. This provides the structural information to form an image. The exit wave immediately below the sample, 𝜓, depends on the probe position *r*_*i*_ and the relative position *r* in a plane perpendicular to the optical axis. For coherent as well as partially coherent modes in mixed-state conditions, the waves are added as complex numbers *Ae*^*iφ*^. The intensity (count of electrons) measured in the diffraction plane at location *k* = (2*π*/*λ*)*θ* is expected as

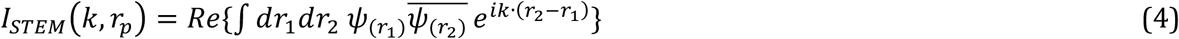

The notation *Re* means that only the real part contributes to intensity. We interpret Eq. 4 as the sum of interfered exit waves, as shown in Fig.S9 (strictly the relation to interference comes from 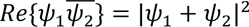 − background). We further limit ourselves to the kinematic approximation, meaning that we ignore multiple scattering, i.e., the sample thickness is not much larger than the mean free path. Thus, effective contributions of interfered waves contain only a phase difference Δ*φ* due to a feature in relation to the average phase delay of the sample reflecting the mean inner potential. There is also a phase difference due to optical aberration in the illumination according to the path difference Δ*χ*. Thus

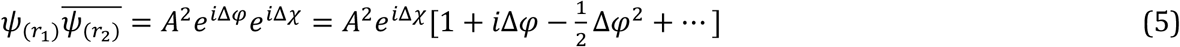

The result is an expansion in orders of the phase shift convolved with order-specific contrast transfer functions

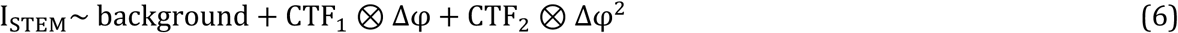

The same expansion is easily obtained for TEM based on Frank’s 1973 work (*51*), and this similarity is expected based on reciprocity in the optical direction. Without aberration, Δ*χ* = 0, the first-order term in Eq.5 is purely imaginary and does not generate contrast. The second-order term is called amplitude contrast. It emerges in the absence of aberration (i.e., at focus) and has the effect of an attenuation of the transmitted signal integrated over the incident beam range in *k*. For a non-zero aberration, which is in general a function of *k*, familiar phase contrast is generated by the cross term *i*Δχ(*k*) *i*Δ*ϕ*(*r*).

The form of Eq.6 applies also to iDPC, and the details can be found in Lazic & Bosch (*27*). In practice, iDPC is normally integrated from difference signals on quadrants of an azimuthally segmented detector. Each detector element generates a complete image after scanning; by reciprocity, the four images are similar to TEM images acquired with tilted illumination. In this case, a non-zero defocus aberration results in image shifts that can be recognized as a parallax effect.

The calculation of parallax corrected iDPC involves deshifting (translating) the images I1, I2, I3, I4 compared to one another by the amount Δ*l* (see Fig.1D), and finding the DPC vector field from the two images

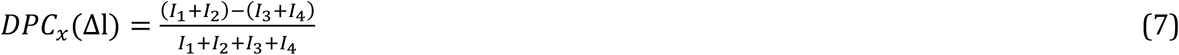

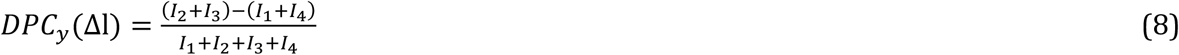

followed by integration

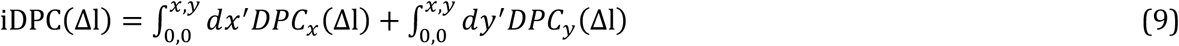

The effect of parallax correction is to reconstruct, at least approximately, the iDPC image that would have been acquired in focus. By comparison with Eq.6, the difference between the original iDPC and the parallax-corrected iDPC provides the amplitude term proportional to the defocus CTF_2_ ⊗ Δ*φ*^2^. Accordingly, we showed (*29*, Appendix B) that at low spatial frequencies CTF_2_ is proportional to defocus. When the defocus is small, however, this term is not useful. As explained in the following, however, we can exploit the linearity of CTF_2_ to remove this constraint.

The parallax differentiated iDPC (piDPC, or πDPC) is defined formally as

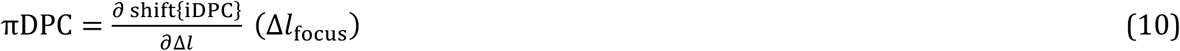

where Δ*l*_focus_ is as above, the image shift determined to compensate for the defocus aberration. The practical calculation is done discretely:

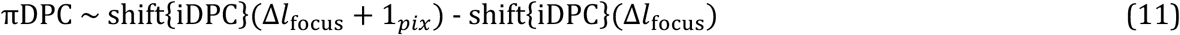

The result is again related to the second-order contrast term

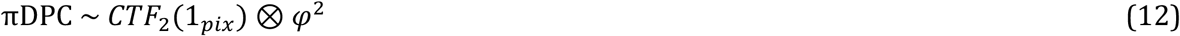

but is not critically dependent on the acquisition defocus.

Note that for a given parallax shift Δ*l*_focus_, an additional defocus of one depth of field unit *λ*/*α*^2^ corresponds to the variation 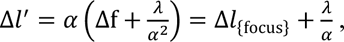 where Δf is the defocus and *α* is the illumination semi-convergence angle. As the pixel sampling is normally chosen as a fraction of the lateral resolution scale *λ*/*α*, so the additional shift of 1 pixel represents the same fraction of the depth of field unit.

In thick samples, we find that the πDPC provides consistently excellent contrast. The simple parallax-corrected iDPC image is clear and sharp for thin specimens such as nanotubes, but is typically poor in a tomographic reconstruction of a thick sample. Apparently, the underlying assumption of a linear relation between intensity and accumulated phase delays is violated after parallax is taken into account. Indeed, thick samples such as cells do not conform to the weak phase object approximation, so the πDPC offers a means of analysis suitable for extension to thick specimens.

### Data processing

For data processing, a Python script cycles through all the data folders, executing a Matlab script in each data folders which performs the deshifting of the individual segments and iDPC analysis (see supplementary information and GitHub). The Matlab script writes out the assembled pBF, iDPC, iDPC1, iDPC2, πDPC, and sBF tilt series. Tilt series alignment was performed using standard methods, in this case using IMOD. Alignment has been performed on the tilt series with the best contrast and then transferred to the other tilt series of this dataset using a short bash script. Subsequent 3D-deconvolution (*32*) is performed as described in Kirchweger et al. (*33*) and Seifer et al. (*23*) by executing another Python script. All scripts are available on GitHub.

### Segmentation

Segmentation and 3D representation of the data were performed using Amira® 2024.2 (Thermo Fisher Scientific, USA). Most structures were segmented manually, based on contrast differences and the characteristic shapes of each component, while collagen fibers were segmented mostly automatically using their distinctive contrast. The resulting 3D structures were visualized using the Generate Surface module.

The 3D orientation of fibers in Amira was calculated using the Label Analysis module. Fibers were manually separated into individual materials, and the software treated each material as a cloud of voxels, computing its principal axes from the voxel distribution to represent the main directions of elongation. The longest principal axis, corresponding to the direction in which the fiber is most stretched, is defined as the fiber’s orientation and reported as an angle relative to the Z-axis.

## Author contributions

Conceptualization: P.K., S.S., B.Z., F.K.M.S., M.E.

Methodology: P.K., S.S., N.V., B.Z., F.K.M.S., M.E.

Investigation: P.K., B.Z.,

Visualization: P.K., N.V.

Supervision: S.G.W., F.K.M.S., M.E.

Writing—original draft: P.K., S.S., M.E., B.Z.

Writing—review & editing: P.K., S.S., B.Z., F.K.M.S., M.E.

## Data and materials availability

The πDPC tomograms have been submitted to the electron microscopy databank with the following EMDB IDs: EMD-55812 (Fig. 4), EMD-55810 (Fig. 5), and EMD-55811 (Fig. 6). The scripts for azimuthal segment data collection from the Nexperion Serial EM script repository (https://serialemscripts.nexperion.net/script/85). The xiDPC repository on GitHub (https://github.com/elbaum-lab/xiDPC-STEM) contains all the scripts for data collection and analysis.

