## Supplemental Material for "Azimuthal Segment Imaging in cryo-STEM Tomography"

**Affiliations**
1 Department of Chemical and Biological Physics, 2 Department of Chemical Research Support, Weizmann Institute of Sciences, Rehovot, Israel 3 Institute of Science and Technology Austria (ISTA), Klosterneuburg, Austria

**This PDF file includes:**

Supplementary Text
Figs. S1 to S8
Movies S1 to S5

**Other Supplementary Materials for this manuscript include the following:**

Movies S1 to S5
Tilt series, tomograms and supplementary movies are available on Zenodo: Fig. 4: doi:10.5281/zenodo.17540667; Fig 5: doi:10.5281/zenodo.17587490; Fig 6: doi:10.5281/zenodo.17579990. The scritps for azimuthal segment data collection is available from the Nexperion Serial EM script repository and on GitHub. The scripts for pBF and DPC analysis are available on GitHub.

**Supplementary Text**

### piDPC and pBF data collection and processing

#### Setup

1. Install AutoScript TEM on the microscope computer and on the SerialEM computer
2. Make sure that AutoScript TEM on the microscope computer is reachable from SerialEM:
2.1 Set the correct IP adress or computername
2.2 Test by e.g. running the '20250609_PreAquAtItems.txt' script
3. Center the beam on the detector (e.g. by running the '12segments.py' on the microscope computer, or the 20240729_Auto_Diff_shift.txt from SerialEM)

#### Collection of the individual segments of the panther detector using SerialEM

1. Copy the text of the '20250609_PreAquAtItems.txt' and '20250609_Collect_STEM_Segments.txt' to your SerialEM scripts
2. In the 'PreAquAtItems' script set the defocus (usually around -0.5 µm)
3. In the 'Collect_STEM_Segments' set the number of pixels and the dwell time (µs)
4. In the 'AquireAtItems' setup, specify the 'PreAquAtItems' script to run before each item
5. In the Tilt menu, go to 'Run Script in TS', choose the correct script and then 4, to run it before each Record step
6. Specify a 'Dummy Record': in the camera properties set the 'Record mode' to 512 pixels and 0.1 µs dwell time (the lowest possible for each)
7. Set up positions to record tilt series and make sure to save the tilt series into a separate data folder. The 'Collect_STEM_Segments' script will save the data into a subfolder for each item recorded.

#### Data processing

##### Setup Matlab

Requires MatTomo in search path (PEET project: https://bio3d.colorado.edu/imod/matlab.html). Place the 'panther_xiDPC.m' script into the Matlab startup folder.

##### Run the Python script to execute the pBF and piDPC calculation

1. Make the following changes in the xDPC.conf file:
1.1 Specify the Parent directory, i.e. the data folder that contains the subfolders with the data
1.2 Specify the Matlab folder
1.3 Specify the 'search_pattern' (something which makes the script recognizing the file name)
1.4 Specify the 'tilt_step'
1.5 Specify the 'semi_angle'
1.6 Specify the 'wavelength'
1.7 Specify the how far the BF disk extends on the detector, e.g. 'DF_Outer'
2. execute the Process_Pantherdata script using these options:

###### Using config file only

python Process_Pantherdata.py --config xDPC.conf

###### Using command-line arguments only
python Process_Pantherdata.py -d /path/to/data -m /path/to/matlab
###### Mix config file with command-line overrides
python Process_Pantherdata.py --config xDPC.conf --tilt-step 3.0 --semi-angle 1.5

##### Align the resulting tilt series of a contrast mode (either pBF or piDPC) using IMOD or AreTomo
(Please note, the 'Dummy Record Tilt series' can be used to extract the .rawtlt)
Make sure that AreTomo exports the IMOD files

##### Apply the alignment to other contrast modes
1. Create a new folder
2. Link the relevant tilt series into the new folder
2.1 Merge the tilt series of a contrast mode, e.g. using the IMOD program 'clip add', if they contain useful signal, e.g. inner + middle + outer ring
3. Copy/link the .xf, .tlt, tilt.com and newst.com (and the other files specified in the two .com files)
4. Set trimming to yes or no (line 12)
4.1 If to yes, set the amount of z-slices for the final rotated reconstruction (line 13)
5. If needed, change the filename flag (line 86)
6. Run the '20250507_Rings_reconstruction.sh' script which performs the alignment and reconstruction based on the previous alignment

##### Postprocessing using 3d-deconvolution
as described in https://doi.org/10.1016/j.jsb.2023.107982

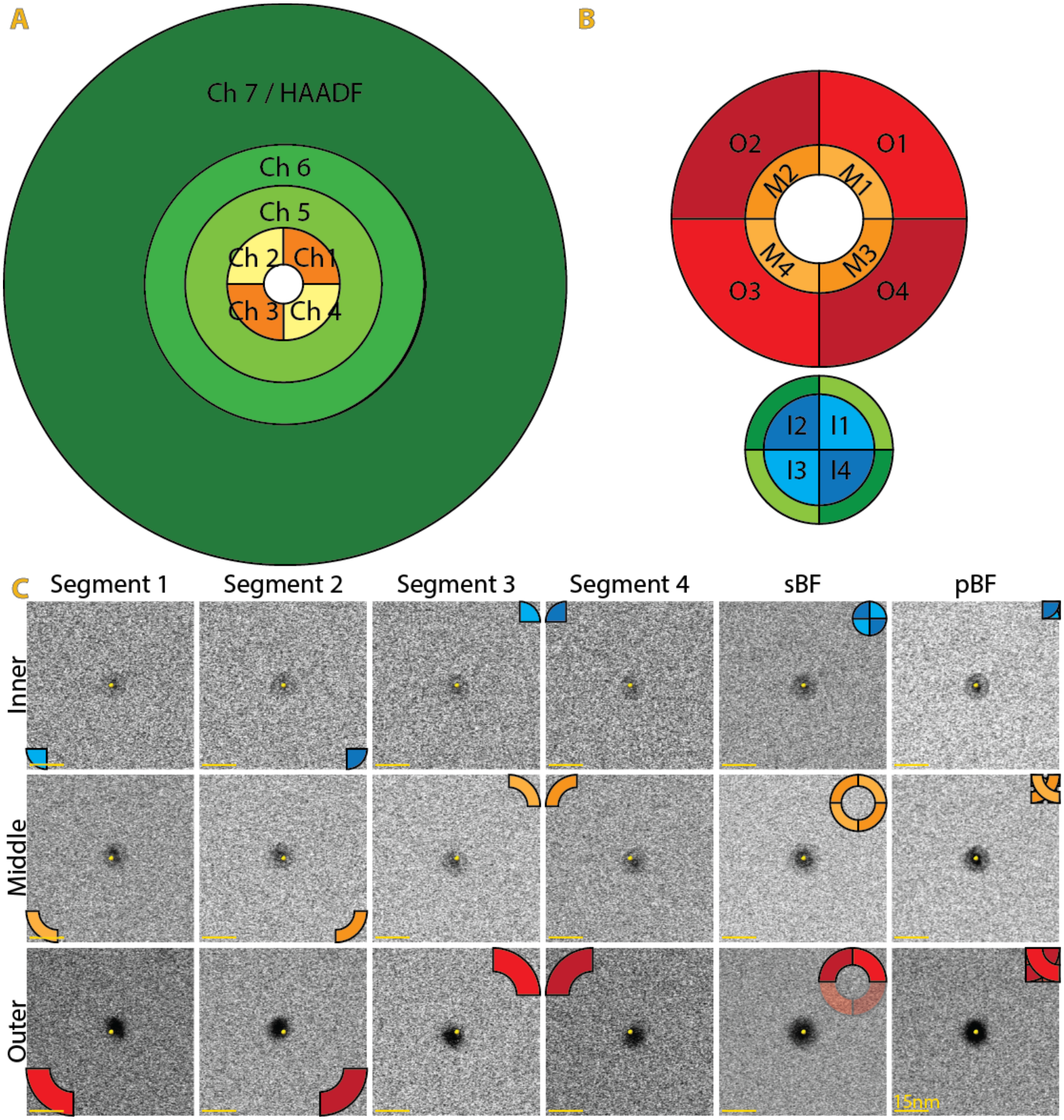


**Fig. S1. Schematic Detector layout (and parallax effect at different collection angles.** Detector layout on the (A) Tecnai T20-F, consisting of the HAADF, and the Opal detector, and (B) Talos Arctica with the Panther detector. (A) The Opal detector consist of an azimuthally segmented ring (Ch1-4, yellow/orange) and two additional rings (Ch5+6). (B) The Panther detector has four rings, all azimuthally segmented, however only the Outer (red), Middle (orange) and Inner rings (blue) are usable simultaneously. (C) Individual (first four columns), summed BF (fifth columns) and pBF (sixth columns) images of a gold bead from the Inner (top row), middle (middle row) and outer (bottom row) rings of the Panther detector.  Yellow dot is in the center of the pBF gold bead. Scale bar is 15 nm.

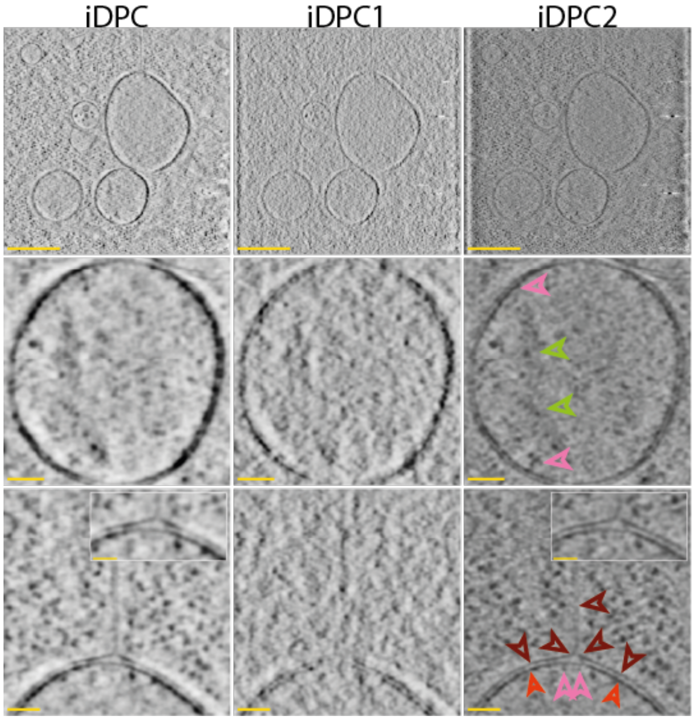


**Fig. S2. Comparison of the iDPC, iDPC1 and iDPC2.** Individual slices of a tomogram recorded on the Opal detector, showing three mitochondrial fragments (top row). Arrowheads and zoom-ins as in Fig 2.

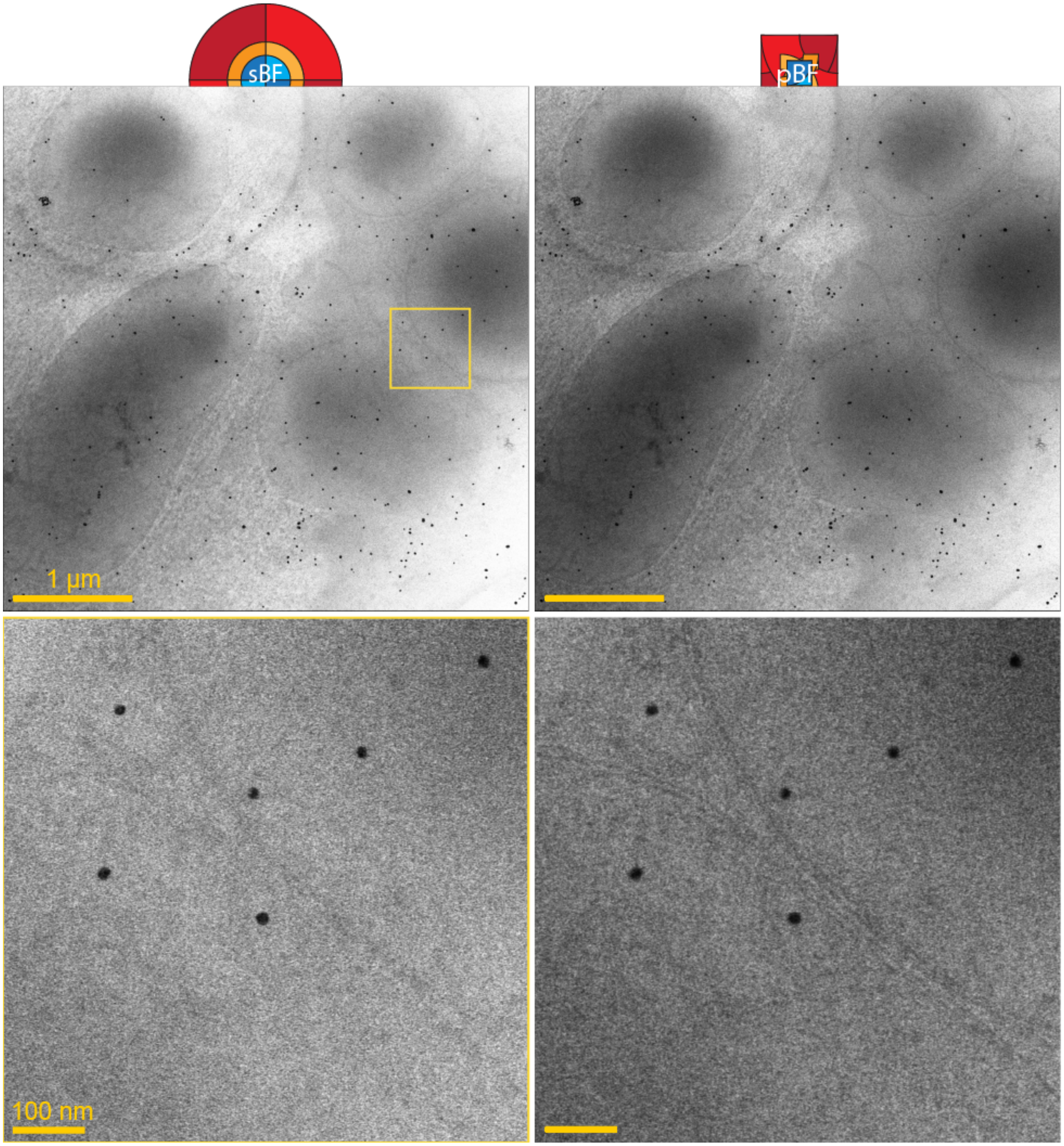


**Fig. S3. Simple BF (sBF) vs pBF.** 0°-tilt of the series that produces the tomogram in Fig 3. Yellow box shows the zoom-in in the bottom row. Scale bars are 1 µm and 100 nm in top and bottom rows, respectively.

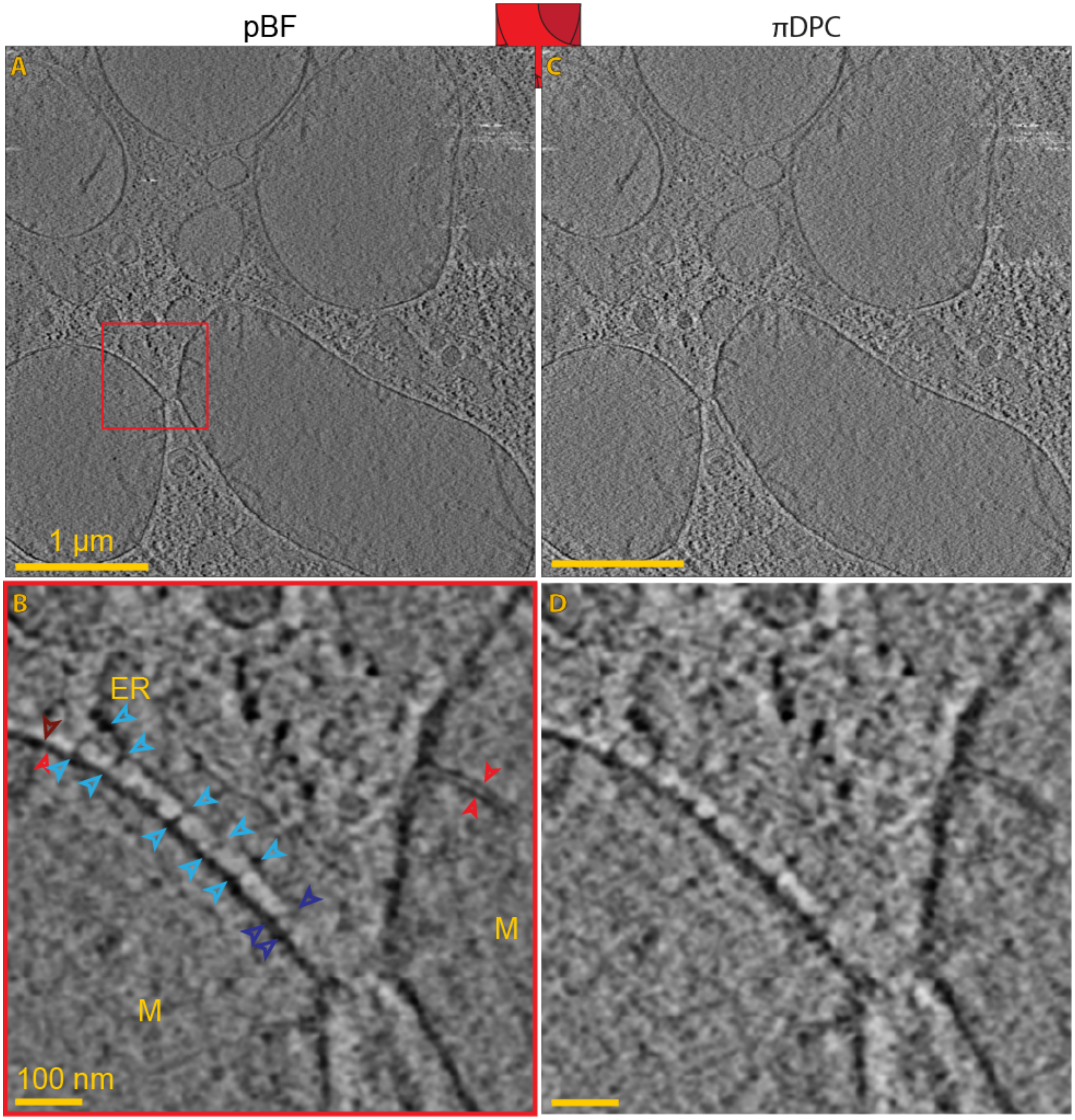


**Fig. S4. Comparison of the pBF and the πDPC from the outer ring.** Description as in Fig 3: Comparison of the pBF with πDPC from the outer ring. Scale bars are 1 µm and 100 nm in the top and bottom rows, respectively.

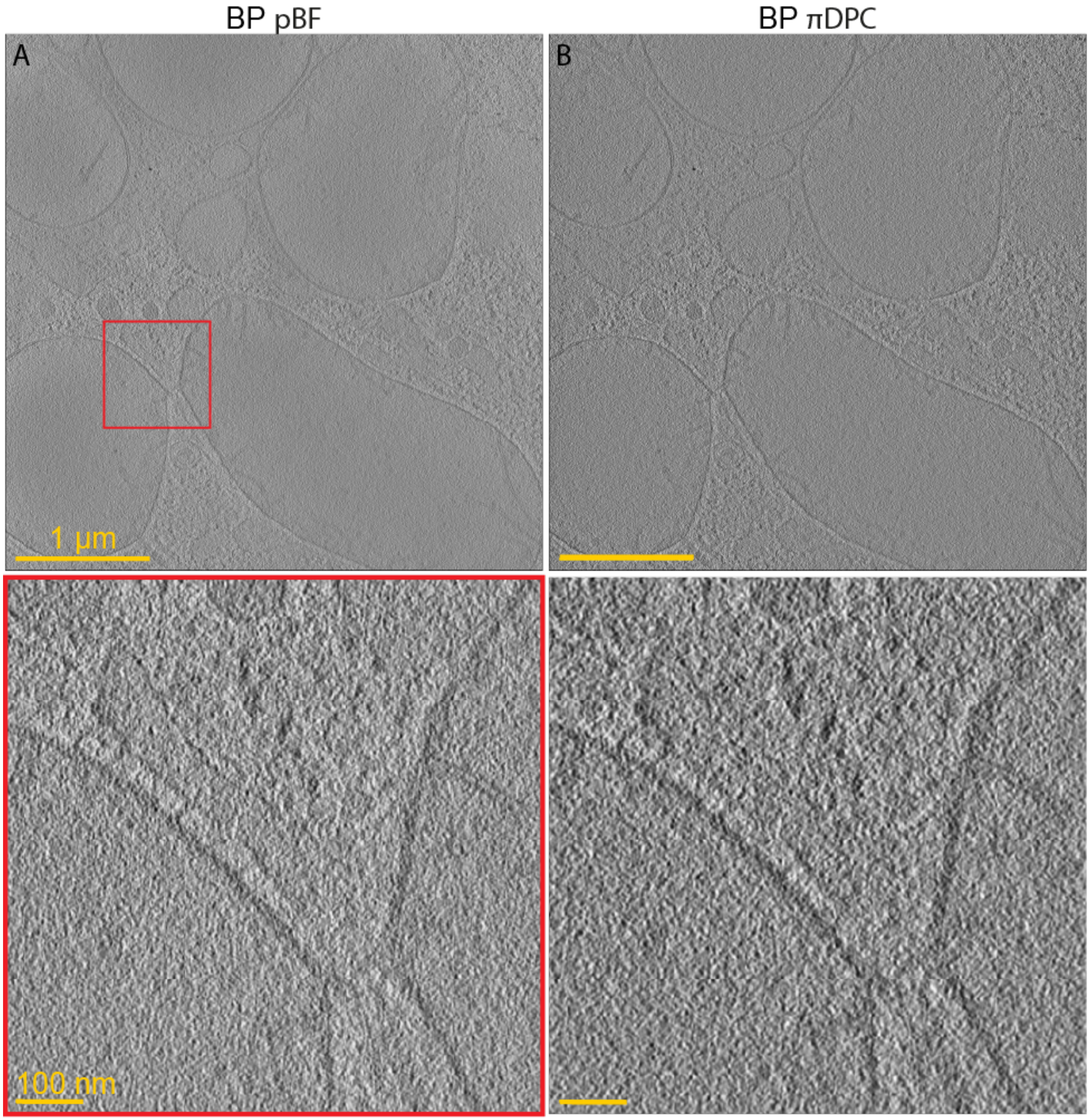


**Fig. S5. Comparison of the back projection of the pBF and the πDPC.** Description as in Fig 3: WBP of the pBF with πDPC from the summed detector. Scale bars are 1 µm and 100 nm in the top and bottom rows, respectively.

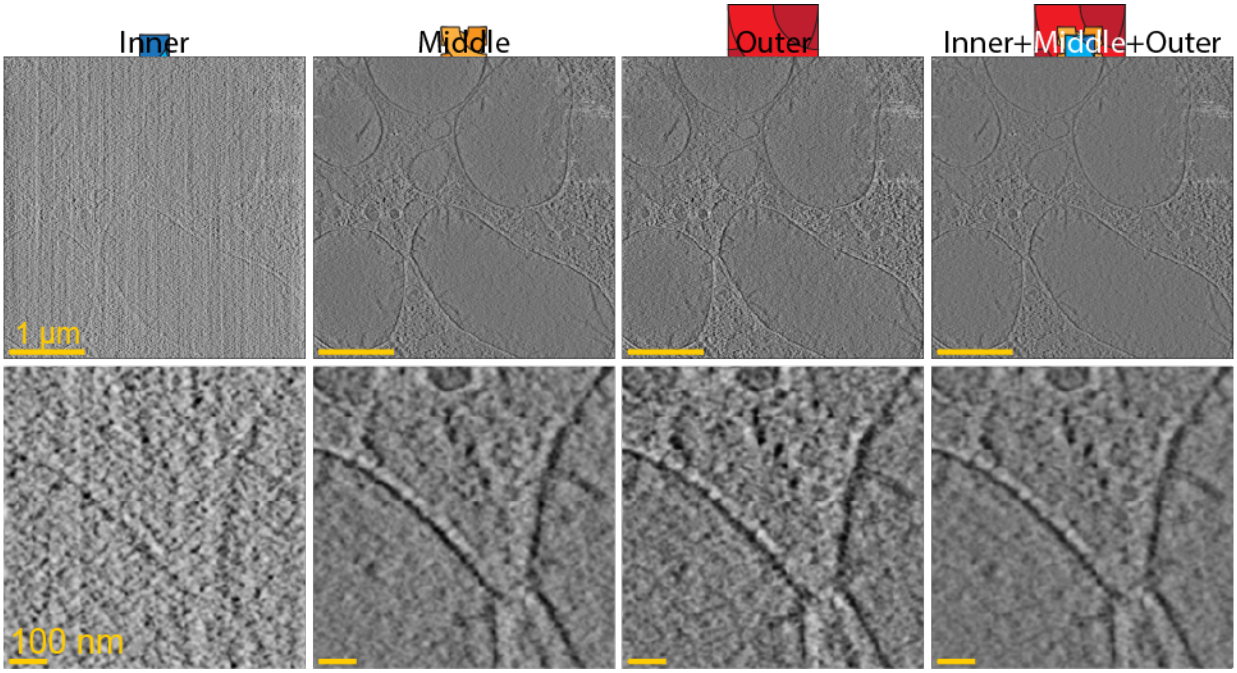


**Fig. S6. Comparison of the πDPC of the inner, middle and outer ring, and the three rings merged into one.** Description as in Fig 3. Same description as in Fig 3. Outer is the same images as in Fig S4B. Inner+Middle+Outer is the same image as in Fig 3B. Scale bars are 1 µm and 100 nm in the top and bottom rows, respectively.

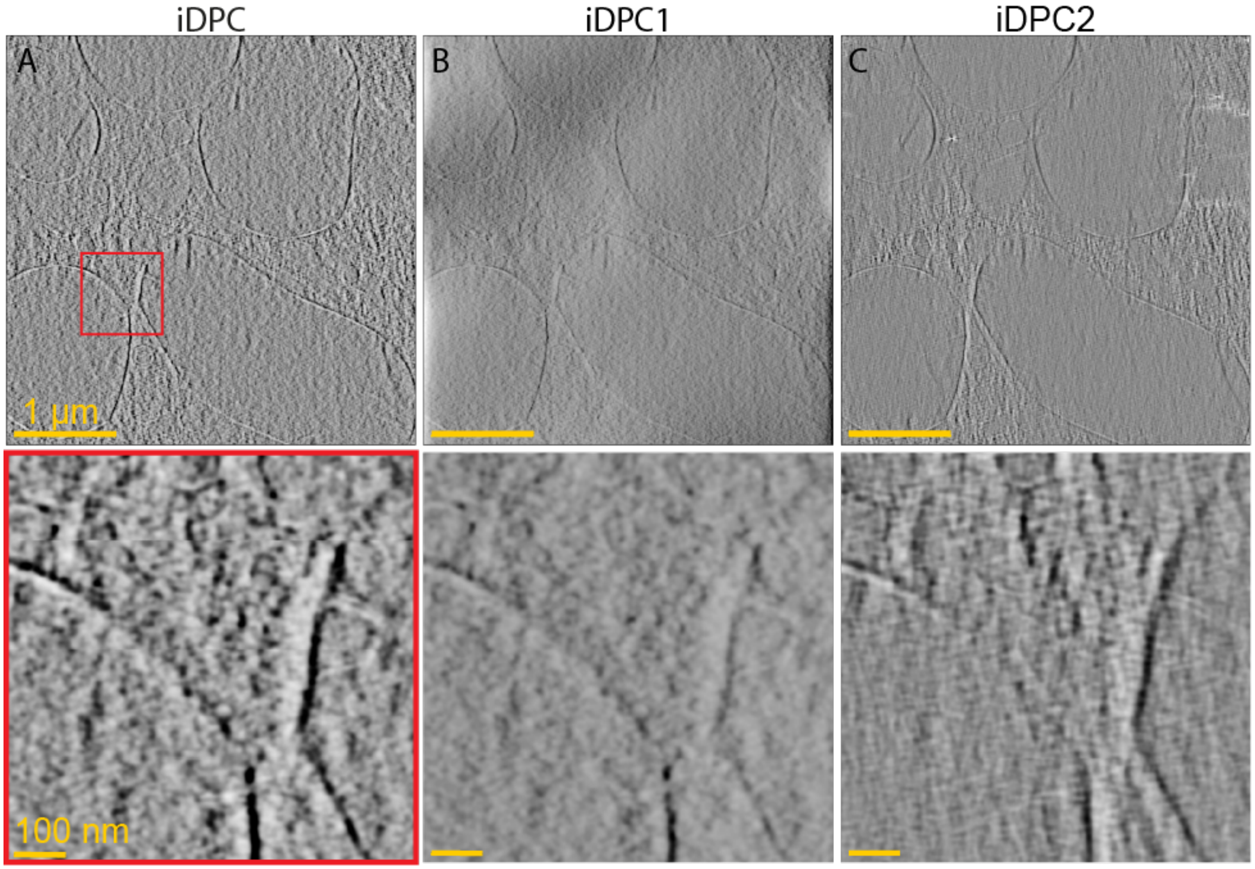


**Fig. S7. Comparison of the deconvolution of iDPC, iDPC1, and iDPC2.** Description as in Fig 3. Scale bars are 1 µm and 100 nm in the top and bottom rows, respectively.

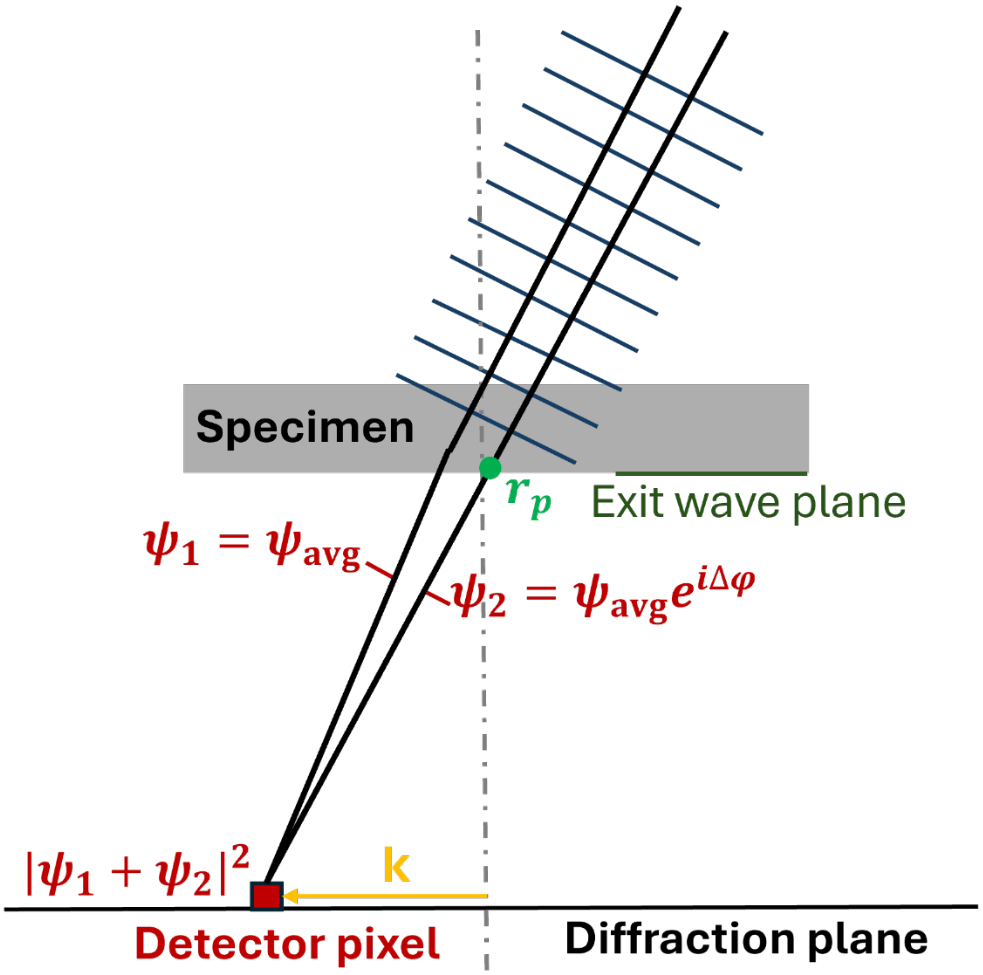


**Fig S8:** The intensity at the detector pixel is viewed as interference between a wave scattered by a phase object at the probe position $r_p$ and many scattered waves that average to the general composition of the specimen.

**Movie S1.**
Individual images of the outer ring of the Panther detector of a gold bead (yellow dot is in the center of the pBF gold bead). Scale Bar is 15 nm.

**Movie S2.**
Individual images of the outer ring of the Panther detector of a T4 bacteriophage. Scale Bar is 50 nm.

**Movie S3.**
πDPC imaging of the CDMs from the Panther detector. The movie shows slices through the tomogram plus the segmentation. Same as in Fig 5.

**Movie S4.**
πDPC imaging of the CDMs from the Panther detector. The movie shows slices through the tomogram plus the segmentation. Same as in Fig 6.

**Movie S5.**
πDPC imaging of the CDMs from the Panther detector. Highlight of the collagen fibers in group 3 and 4. Same as in Fig 7C. Scale bar is 500 nm.
